# Specialization of mid-tier stages of dorsal and ventral pathways in stereoscopic processing

**DOI:** 10.1101/2020.03.10.985721

**Authors:** Toshihide W. Yoshioka, Takahiro Doi, Mohammad Abdolrahmani, Ichiro Fujita

**Affiliations:** Laboratory for Cognitive Neuroscience, Graduate School of Frontier Biosciences, 1–4 Yamadaoka, Suita, Osaka 565–0871, Japan; Center for Information and Neural Networks, Osaka University and National Institute of Information and Communications Technology, 1–4 Yamadaoka, Suita, Osaka 565–0871, Japan; Department of Psychology, University of Pennsylvania, Philadelphia, PA 19104, USA; Laboratory for Neural Circuits and Behavior, Center for Brain Science (CBS), 2-1 Hirosawa, Wako, Saitama, 351-0198, Japan

**Keywords:** stereopsis, binocular vision, extrastriate visual cortex, disparity energy model, false-match rejection, macaque

## Abstract

The division of labor between the dorsal and ventral visual pathways is an influential model of parallel information processing in the cerebral cortex. However, direct comparison of the two pathways at the single-neuron resolution has been scarce. Here we compare how MT and V4, mid-tier areas of the two pathways in the monkey, process binocular disparity, a powerful cue for depth perception and visually guided actions. We report a novel tradeoff where MT neurons transmit disparity signals quickly and robustly, whereas V4 neurons markedly transform the nature of the signals with extra time to solve the stereo correspondence problem. Therefore, signaling speed and robustness are traded for computational complexity. The key factor in this tradeoff was the shape of disparity tuning: V4 neurons had more even-symmetric tuning than MT neurons. Moreover, this correlation between tuning shape and signal transformation was present across individual neurons within both MT and V4. Overall, our results reveal both distinct signaling advantages and common tuning-curve features of the dorsal and ventral pathways in stereoscopic processing.

## Introduction

The dorsal and ventral pathways of the primate visual system have served as a widely influential model of parallel sensory processing in the mammalian cerebral cortex (Ungerleider and Mishkin, 1982). Similar organizational structures have been found in the visual cortical networks of cats and mice (Hilgetag et al., 2000; Wang et al., 2012). Moreover, related ideas have been put forth regarding non-visual domains such as auditory (Rauschecker and Scott, 2009), somatosensory (Dijkerman and DeHaan, 2007), and language-related processes (Hickok and Poeppel, 2007; Saur et al., 2008). Despite their widespread influence, direct comparison of the dorsal and ventral pathways at the single-neuron resolution is surprisingly scarce (Cheng et al., 1994; Lehky and Sereno, 2007), leaving open the exact nature of the differences and similarities between the two pathways.

In the visual systems of human and non-human primates, the dorsal pathway is implicated in spatial vision and vision for action, whereas the ventral pathway is implicated in object recognition and vision for perception (Goodale and Milner, 1992; Ungerleider and Mishkin, 1982). Contrary to initial speculation, both pathways process the same visual cues. Neurons with shape and color selectivity are found not only in ventral areas but also in dorsal areas (Seidemann et al., 1999; Sereno and Maunsell 1998). Likewise, neurons with motion direction selectivity are found not only in dorsal areas but also in ventral areas (Desimone and Schein, 1987; Li et al., 2013; Mountcastle et al., 1987). Investigating how the dorsal and ventral pathways process the same cue at the single-neuron resolution is critical to better understand the parallel processing strategies in the visual system.

One of the most notable visual cues in this regard is binocular disparity, the positional difference between images projected to the left and right eyes. As a precise cue for stereoscopic depth, binocular disparity is used for many functions, including the recognition of 3D objects, visually guided 3D action (e.g., reaching, grasping, and vergence eye movement), and navigation through the 3D environment (e.g., moving through obstacles). As binocular disparity is used for both “what” (object vision) and “where” (spatial vision) functions, the visual system processes binocular disparity along both the dorsal and ventral pathways (Neri, 2005; Orban et al., 2006; Parker, 2007; Roe et al., 2012; Welchman, 2016).

To encode binocular disparity faithfully in accordance with the geometry of the 3D world, the visual system should identify the corresponding visual features in left and right retinal images that originate from the same surface point in the 3D space (Julesz, 1971; Marr and Poggio, 1979). A conventional method to probe the neuronal process for stereo correspondence is to measure the disparity selectivity for anticorrelated random-dot stereograms (aRDSs) (Chen et al., 2017; Cumming and Parker, 1997; Goncalves and Welchman, 2017; Janssen et al., 2003; Krug et al., 2004; Kumano et al., 2008; Nieder and Wagner, 2001; Ohzawa, 1998; Samonds et al., 2013; Takemura et al., 2001; Tanabe et al., 2004; Theys et al., 2012). In aRDSs, one eye image is contrast-reversed to eliminate the natural solution to the correspondence problem; the natural solution requires the depth of the corresponding points to vary smoothly over space, but such a solution does not exist for aRDSs. Accordingly, the neural correlate of the correspondence solution should have no or minimal disparity selectivity for aRDSs. Although the activity of many V1 neurons encodes binocular disparity, these neurons also falsely encode the disparity embedded in aRDSs (Cumming and Parker, 1997). Simple computations akin to interocular cross-correlation explain this primitive disparity representation (Ohzawa et al., 1990). To solve the correspondence problem, a more complex computation should ensue and transform the primitive correlation-based disparity representation into the sophisticated representation that is based on binocularly matching features (“match-based representation”).

In this study, we directly compared the disparity representations in areas MT and V4 of the monkey. MT and V4 are counterpart mid-tier stages of the dorsal and ventral pathways, both of which causally contribute to stereoscopic depth judgment (DeAngelis et al., 1998; Shiozaki et al., 2012; Uka and DeAngelis, 2006). We used a recently developed extension of aRDSs, called graded anticorrelation (Doi and Fujita, 2014; Doi et al. 2011, 2013). Moreover, we used virtually identical experimental and analysis methods (and one identical animal) in MT and V4 experiments to achieve a more controlled comparison of the two areas than has previously been attempted. We found that the disparity selectivity was indistinguishable between MT and V4 in their responses to the conventional stimuli, aRDSs. However, graded anticorrelation revealed that the disparity representation in V4 was more strongly specialized for the correspondence solution than that in MT. The relative advantage of MT’s disparity signaling was its speed and strength. These results shed light on the division of labor between the dorsal and ventral pathways. The dorsal pathway carries out rapid signaling to produce timely behavioral outputs, whereas the ventral pathway emphasizes complex computations to derive sophisticated sensory representations. We also found that the disparity-tuning shape was highly consistent with the degree of neuronal specialization: neurons with more even-symmetric tuning had responses closer to the correspondence solution both within and across MT and V4. We will discuss this relationship in light of the adaptation of the stereoscopic system to natural binocular inputs (Haefner and Cumming, 2008).

## Results

### Dissociating correlation-based and match-based representations by graded anticorrelation

We analyzed the neuronal responses to graded anticorrelation to characterize how well these responses conformed to the solution of the correspondence problem; that is, how much the disparity tuning of a visual neuron deviated from the simple correlation-based representation toward a more sophisticated match-based representation (Doi and Fujita, 2014; Doi et al. 2011, 2013). For this assessment, we manipulated the binocular correlation of RDSs by reversing the luminance contrast of a varying proportion of dots in one eye (Figure 1A). This stimulus manipulation systematically changed the binocular correlation (correlation between the left-eye and right-eye images; the top arrow in Figure 1A) and the binocular match (the percentage of binocularly matched features; the bottom arrow in Figure 1A) in a dissociable manner. At one end, correlated RDSs (cRDSs) had 100% correlation and 100% match (Figure 1A right). However, at the other end, aRDSs had −100% correlation and 0% match, indicating complete dissociation (Figure 1A left). In the middle, half-matched RDSs (hmRDSs) showed that half of the dots were contrast-reversed in one eye. Thus, the overall binocular correlation of hmRDSs was 0% since correlated and anticorrelated dots canceled out, while the strength (proportion) of the binocular match was 50%, indicating dissociation in the opposite direction.

**Figure 1.**
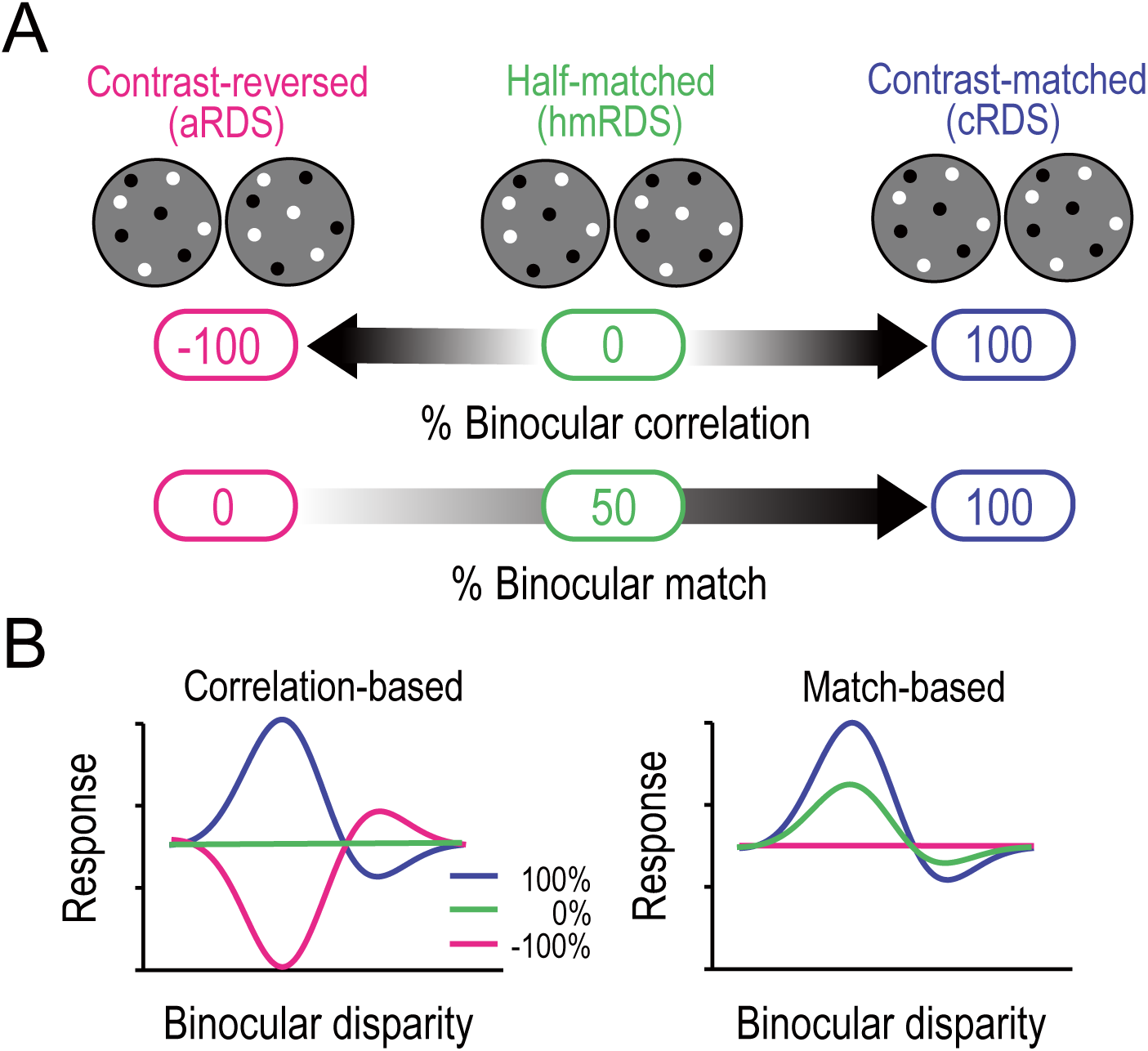
Correlation-based and match-based representations predict distinct responses to graded anticorrelation. (**A**) Three RDSs with graded anticorrelation. From right to left, the fraction of binocularly contrast-matched dots decreases from 100% (correlated RDS or cRDS) through 50% (half-matched RDS or hmRDS) to 0% (anticorrelated RDS or aRDS). This same stimulus manipulation decreases binocular correlation gradually from 100% (cRDS) through 0% (hmRDS) to −100% (aRDS). (**B**) Hypothetical disparity-tuning curves to graded-anticorrelation stimuli as predicted by correlation-based or match-based disparity representations. In the correlation-based responses (**left**), the absolute value and the sign of the percent binocular correlation determine the amplitude and shape of the tuning function, respectively. In the match-based responses (**right**), the amplitude reflects the fraction of matched dots, and the shape is invariant.

We used these dissociated values of binocular correlation and binocular match to derive the predictions of disparity-tuning curves. A disparity detector with the pure correlation-based representation should lose its disparity selectivity and show flat tuning for hmRDS, because hmRDSs offer no binocular correlation that is necessary for these kinds of detectors to encode disparity (Figure 1B left, green line). For aRDSs, the same detectors should have an inverted tuning curve of the same magnitude as the curve for cRDSs, because of the sign inversion of the stimulus correlation (Figure 1B left, pink curve). By contrast, ideal match-based detectors should retain disparity tuning with a decreased amplitude for hmRDSs but should lose disparity selectivity for aRDSs (Figure 1B right). This is because hmRDSs contain binocularly matched features whereas aRDSs offer no correctly matching features between the two eyes (i.e., no natural solution to the correspondence problem). The disparity energy model is a classic implementation of the correlation-based detector (Cumming and Parker, 1997; Ohzawa et al., 1990). Additional nonlinearity can transform the correlation-based detector to the match-based detector under some conditions (Doi and Fujita, 2014; Henriksen et al., 2016a).

### Disparity selectivity in MT and V4 is biased toward correlation-based and match-based representations, respectively

We recorded single-neuron activity from monkeys while they performed a fixation task. The monkey is an excellent model animal for binocular vision because like humans, they have two forward-facing eyes. Moreover, their binocular depth perception is similar to that in humans (Shiozaki et al. 2012; Uka et al., 1999). We compared how neurons in MT and V4 responded to graded anticorrelation by examining the responses of 83 disparity-selective neurons in MT (monkey O, N = 32; monkey A, N = 51) and 78 disparity selective neurons in V4 (monkey O, N = 29; monkey I, N = 49). Disparity selectivity was determined from the responses to cRDSs (Kruskal-Wallis test; *p* < 0.05). Monkey O provided data sets for both MT and V4. We previously analyzed the same data set from V4 (Abdolrahmani et al., 2016), but most of the analyses in the current paper are novel, with two exceptions (i.e., Figures 2B,D for an example neuron and Figure 5E for direct comparison of a conventional metric against our novel metric).

**Figure 2.**
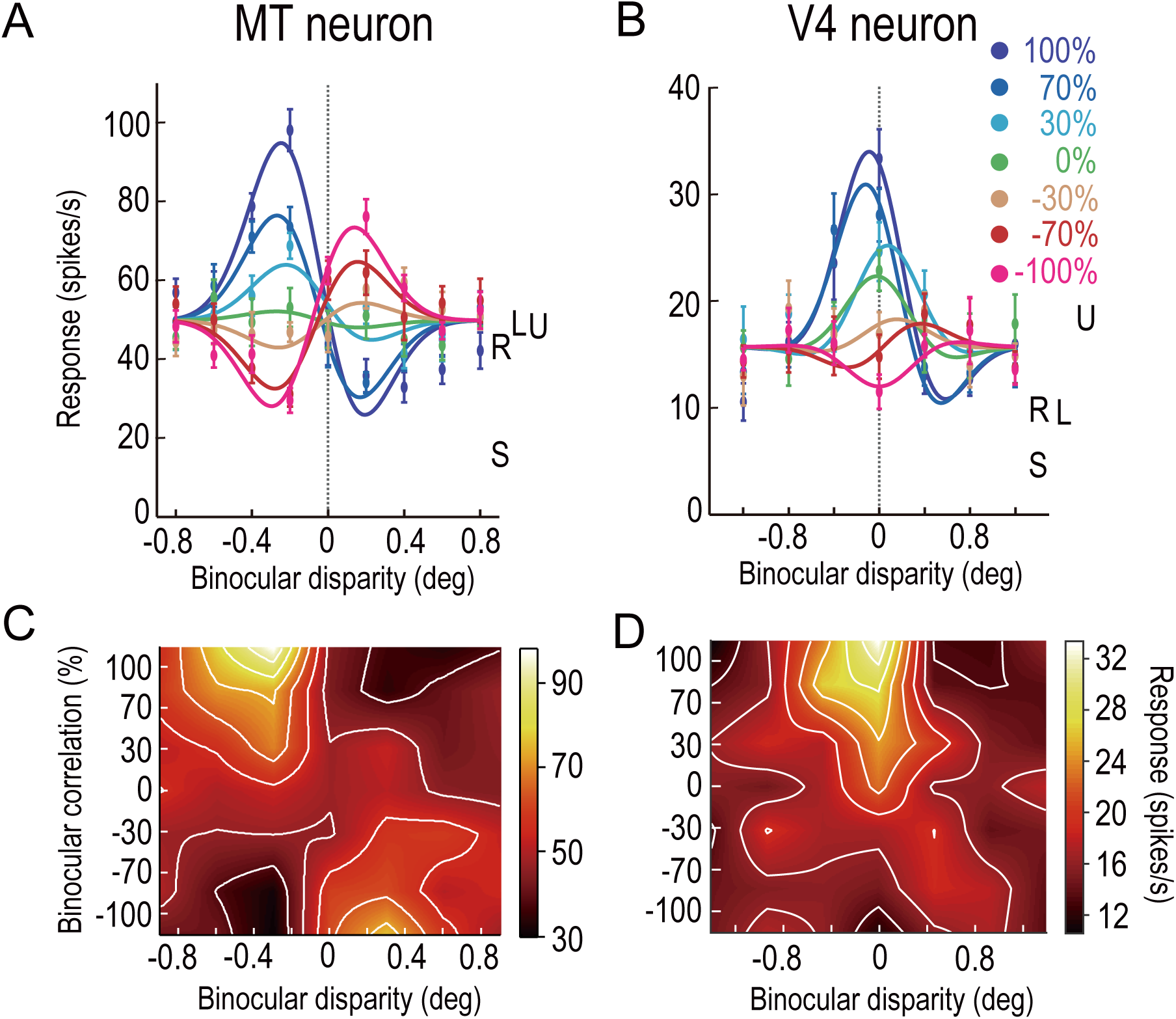
Disparity-tuning curves of example neurons in MT and V4. The average firing rates of example neurons at different binocular-disparity values and binocular-correlation levels in MT (**A**) and V4 (**B**). Different colors indicate different correlation levels. Error bars indicate the SEMs. Letters S, U, R, and L indicate the level of spontaneous (pre-stimulus) activity, the response to uncorrelated RDSs, and the responses to right and left monocular images, respectively. (**C, D**) Color plots of responses of the neurons shown in **A** and **B** in a 2D plane defined by binocular disparity and correlation level. Brighter colors indicate stronger responses, as shown in the scale bars. The raw responses are linearly interpolated in both dimensions. The V4 example was previously shown in Abdolrahmani et al., 2016.

**Figure 5.**
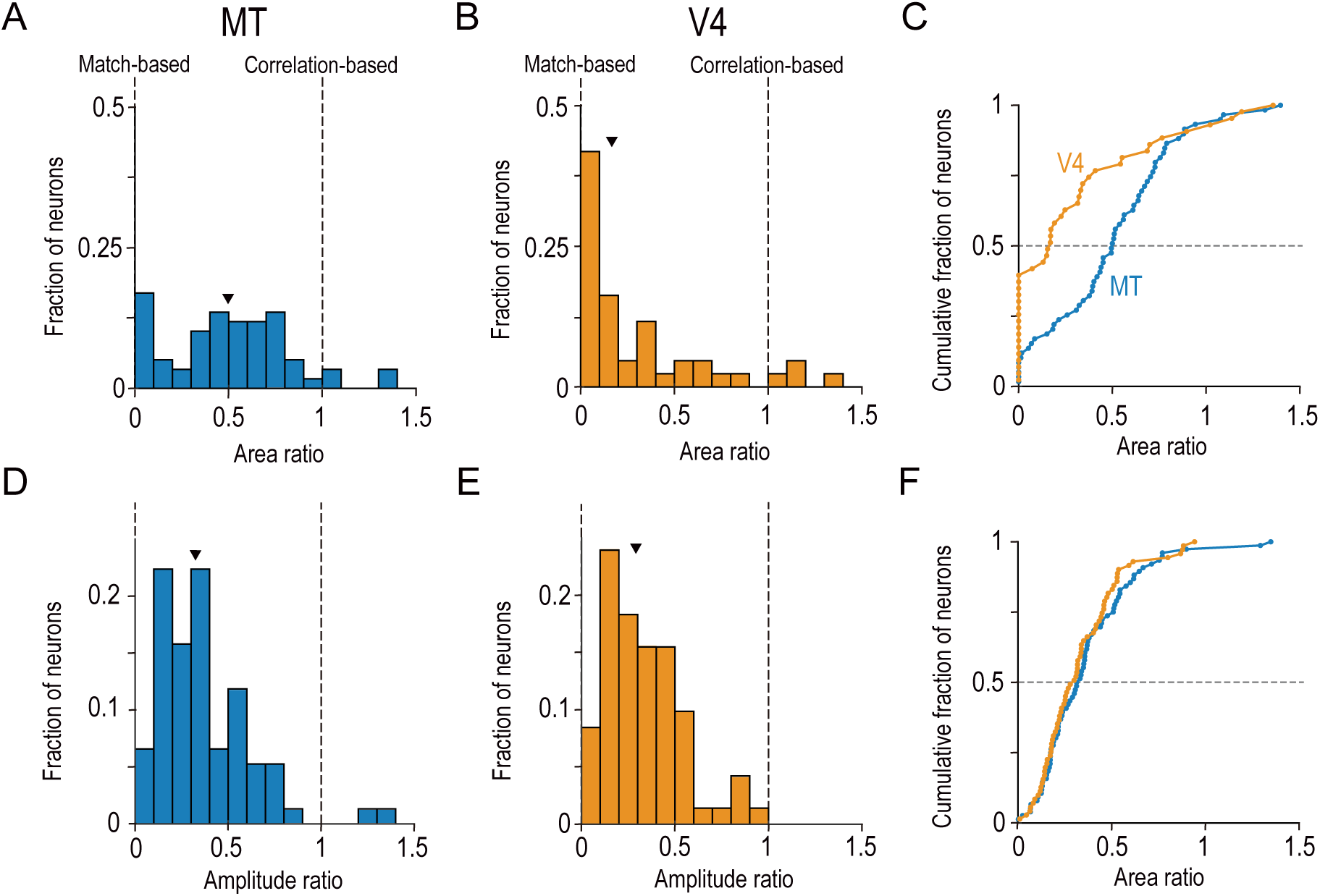
Distinct distributions of single-neuron disparity representation between MT and V4 revealed only by area ratio. Distributions of the area ratio, our new metric (**A–C**), and the amplitude ratio, the conventional metric (**D–F**), are shown. **A–C** consist of the neurons with good Gabor-function fitting to disparity-tuning curves and good quadratic-function fitting to signed amplitude ratios (R^2^ > 0.6; MT, N = 59; V4, N = 43). **D–F** consist of the neurons with good Gabor-function fitting (R^2^ > 0.6; MT, N = 76; V4, N = 71). Triangles indicate the medians. **C** and **F** show cumulative distributions of the area ratio and amplitude ratio, respectively. The amplitude ratio histogram for V4 (**E**) is a replot of the result reported in Abdolrahmani et al., 2016.

The graded anticorrelation of RDSs differentially modulated the disparity tuning of neurons in MT and V4. Figure 2 shows example single neurons demonstrating characteristic differences between the MT and V4 populations. Both of the example neurons responded maximally at “near” disparities with 100% binocular correlation, but the decrease in the correlation level changed the tuning curves differentially between them. The change observed for the example MT neuron is reminiscent of the correlation-based representation of disparity: when the correlation level decreased from 100% to 0% (hmRDS), the tuning amplitude gradually decreased to almost zero (flat curve). A further decrease in the correlation level from 0% to −100% (aRDSs) recovered the tuning but with an inverted shape (Figure 2A). Compared to this MT example, the response pattern of the example V4 neuron was closer to the match-based representation: the tuning amplitude was noticeably retained at 0% correlation with a relatively unchanged tuning phase (shape). The tuning was more attenuated than the MT example at −100% correlation, without the exact shape inversion (Figure 2B). We plotted these responses in the 2D plane defined by binocular disparity and correlation level. The same pattern was also apparent when we compared the observed 2D response profiles (Figures 2C,D) to the predicted profiles from the correlation-based and match-based representations (Figure S1). Below we quantify and validate this impression with both model-free and model-based analyses

The disparity representation was drastically different between neurons in MT and V4 at the population level. MT neurons retained the characteristics of the correlation-based disparity representation presumably constructed in V1. These features were largely eliminated in V4, where the disparity selectivity was strongly biased toward the match-based representation. Figure 3 shows the results of model-free population analyses without curve fitting. First, we analyzed the fraction of disparity-selective neurons in MT (Kruskal-Wallis test, *p* < 0.05 tested independently at each correlation level below 100%). The fraction had a characteristic U shape consistent with the correlation-based representation: disparity encoding relies on the non-zero binocular correlation of the stereo images (Figure 3A). The fraction was minimal at −30% correlation, where only one out of 83 MT neurons (1.2%) was disparity selective. With further decrease from −30% to −100%, the fraction increased to 21/83 (25.3%). The pure correlation-based model posits that the fraction should be 1 at −100% correlation and the minimum at 0% correlation. The mismatch slightly hints at the match-based representation, although overall the disparity selectivity in MT is correlation-based.

**Figure 3.**
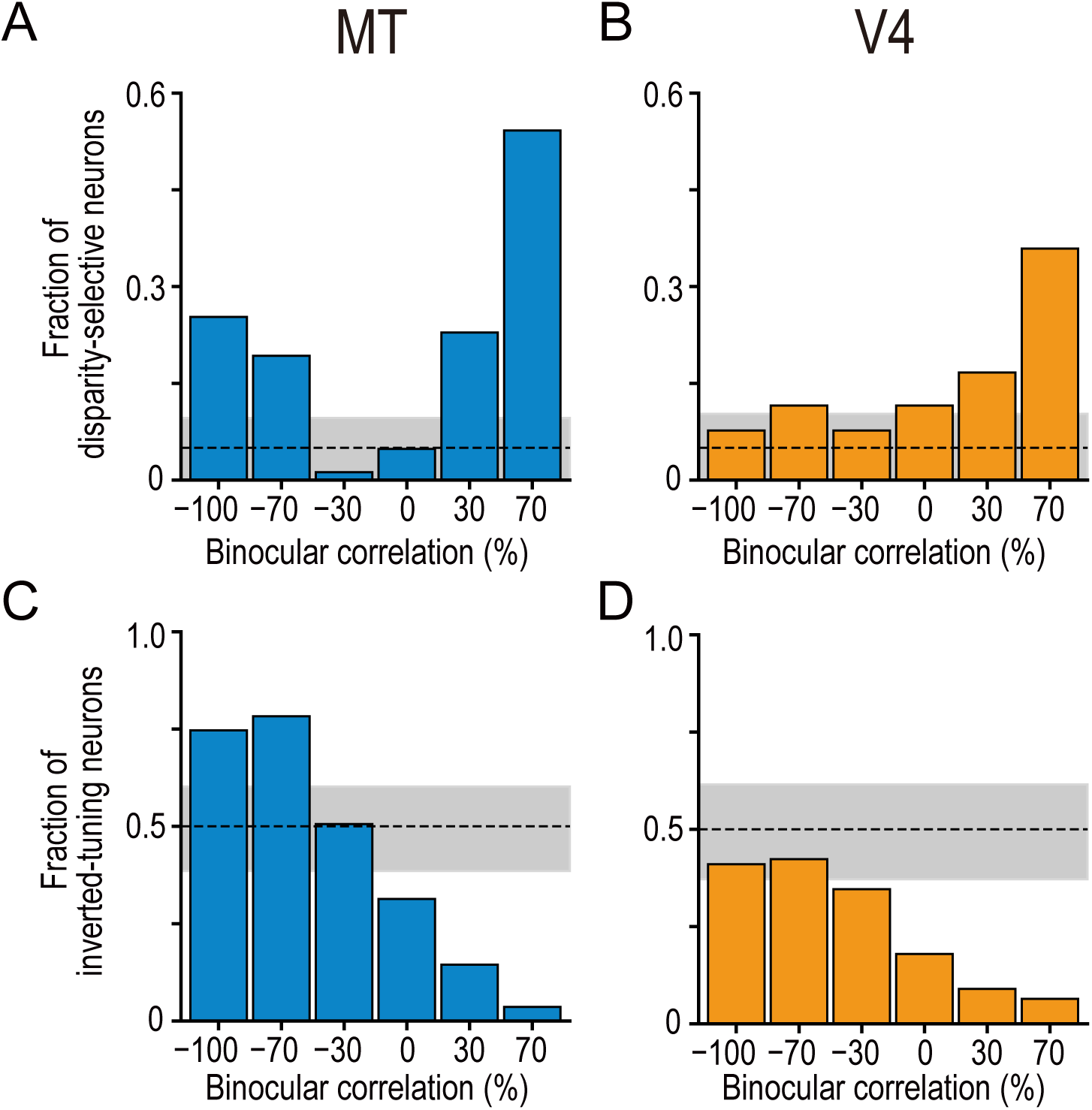
Fractions of disparity-selective neurons and inverted-tuning neurons change with graded anticorrelation differently between MT and V4. (**A, B**) The fraction of disparity-selective neurons as a function of binocular correlation. The dashed line indicates the chance level (0.05) and the gray area indicates its 95% interval based on the binomial distribution. The fraction at 100% correlation is one because of our cell selection procedure. (**C, D**) The fraction of neurons with inverted disparity-tuning (relative to the tuning to cRDSs) as a function of binocular correlation. The chance level is 0.5. The fraction at 100% correlation is zero by definition. The figure is based on all the disparity-selective neurons recorded (as determined by the responses to cRDSs; 83 and 78 neurons for MT and V4, respectively).

The tuning shape of MT neurons also exhibited the characteristic of the correlation-based representation: negative binocular correlation inverts the tuning shape (i.e., the preferred disparity for positively correlated RDSs becomes the null disparity for negatively correlated RDSs and *vice versa*). We detected tuning inversion based on a negative correlation coefficient between the responses to cRDSs and those at a lower binocular correlation level. In MT, the fraction of the neurons with inverted tuning gradually increased as binocular correlation decreased (Figure 3C). Notably, significant fractions of neurons inverted their tuning curves at the two most negative correlation levels we tested (75% [62/83] and 78% [65/83] of neurons at the −100% and −70% correlation levels, respectively; *p* = 3.8×10^−6^ and 1.1×10^−7^; H0: the fraction is 0.5; binomial test). We also observed some deviation from the pure correlation representation: the fraction of the tuning inversion was not 1 at the −100% correlation level. The fraction was closest to 0.5 at the −30% correlation level, echoing the finding that the fraction of disparity-selective neurons was minimal at −30% (Figure 3A).

The V4 population data were markedly different from the MT data, entirely lacking the aforementioned characteristics of the correlation-based representation: the fraction of disparity-selective neurons did not have the characteristic U shape. Less than half of the neurons showed the tuning curve inversion at any correlation levels tested. Instead, the disparity selectivity of V4 neurons was mostly consistent with the match-based representation. With gradual anticorrelation, the fraction of disparity-selective neurons gradually decreased (Figure 3B), mirroring the decrease in the level of the binocular match (Figure 1A). This implies that the disparity selectivity in V4 mainly depends on the strength of the binocular match (i.e., the proportion of dots with binocularly matched contrasts). The gradual increase in the fraction of inverted tuning (Figure 3D) is also consistent with the match-based representation, because when the tuning curves become flatter at lower correlation levels, more neurons can be taken to have inverted tuning due to response noise (Figure 1B, right). Just as the MT data were not completely consistent with the pure correlation-based representation, the V4 data were not consistent with the pure match-based representation. See the Discussion section for how aggregating the population responses in V4 would help create the pure match-based representation.

The previous analysis averaged the response characteristics of individual neurons within MT and V4 at each correlation level. Next, we characterized how each individual neuron represented binocular disparity. To this end, we extended a frequently used metric, the amplitude ratio, which quantifies the disparity-tuning amplitude at −100% correlation (aRDS) relative to that at 100% correlation (cRDS; Cumming and Parker 1997; Krug et al., 2004; Kumano et al., 2008; Samonds et al., 2013; Tanabe et al., 2004). First, we added a sign to the amplitude ratio, where a negative sign indicates tuning inversion. Second, we calculated the signed amplitude ratio for all intermediate correlation levels, not just for −100%. This signed amplitude ratio function should have different shapes for the correlation-based and match-based neurons (Figure 4A). For the correlation-based neurons, it should decrease from 1 through 0 to −1 as binocular correlation decreases from 100% through 0% to −100%. For the match-based neurons, it should decrease as the correlation level decreases but should always stay above 0 except at −100% correlation. We do not argue that the match-based prediction must be exactly linear as shown in Figure 4A; the linear decrease is just the simplest example of monotonic functions.

**Figure 4.**
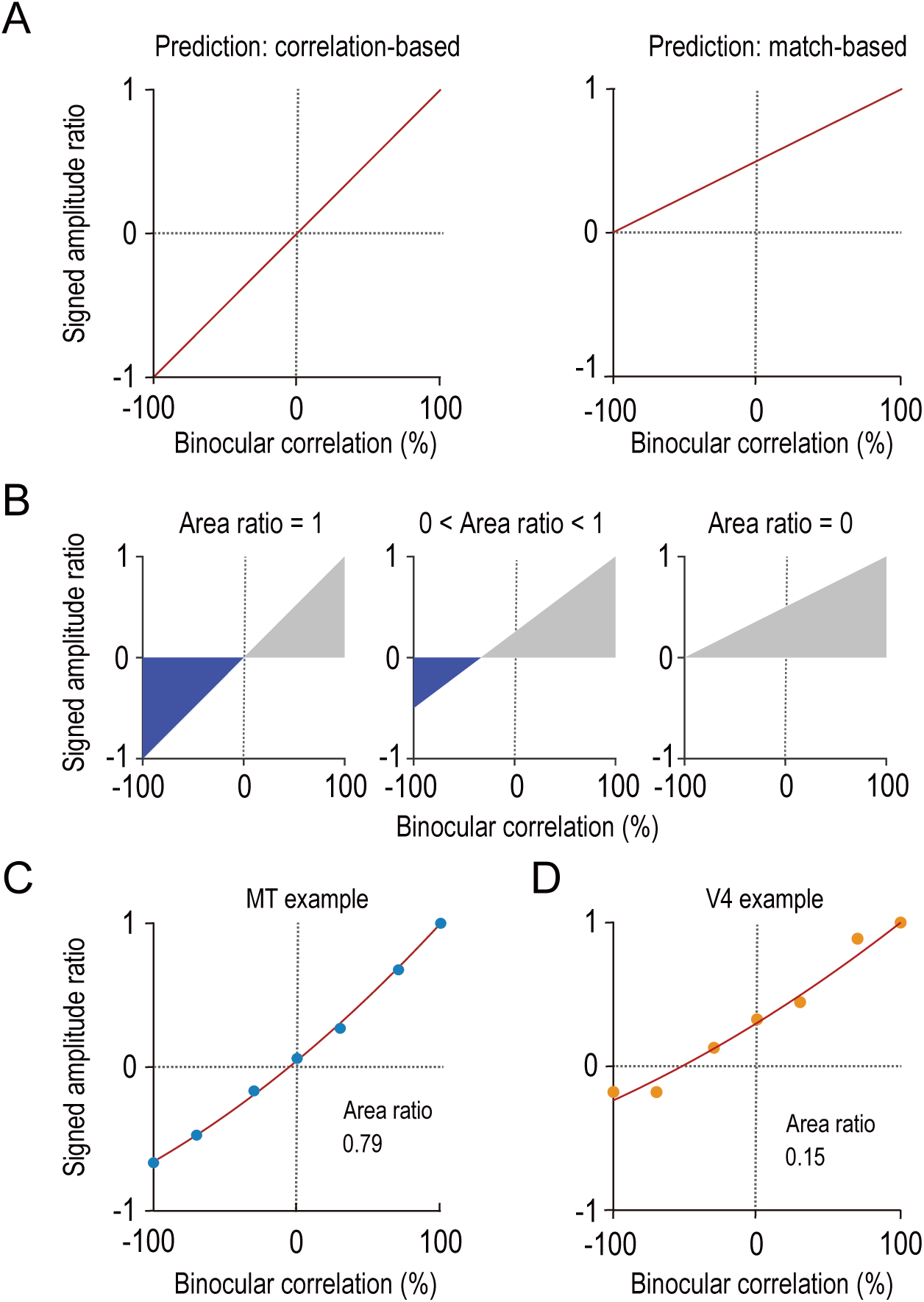
Characterizing disparity representation for each individual neuron. (**A**) Correlation-based and match-based disparity representations predict different profiles for the signed amplitude ratio as a function of binocular correlation. The correlation*-*based representation predicts a monotonic decrease of the signed amplitude ratio from 1 for cRDSs through 0 for hmRDSs to −1 for aRDSs (***left***), whereas the match-based representation predicts a monotonic decrease from 1 for cRDSs through an intermediate positive value for hmRDSs to 0 for aRDSs (***right***). (**B**) Area ratio as a metric to quantify how well the observed responses conform to correlation-based or match-based prediction. The area ratio was defined as the ratio of two areas: the numerator was the area with the negatively signed amplitude ratios (**blue**); the denominator was the area with the positively signed amplitude ratios (**gray**). An area ratio of one indicates perfectly correlation-based responses (***leftmost***), and an area ratio of zero indicates perfectly match-based responses (***rightmost***). An intermediate area ratio indicates an intermediate representation type (***middle***). (**C, D**) Area ratios calculated for the example MT and V4 neurons shown in Figure 2. The observed signed amplitude ratios were interpolated with best-fitted quadratic functions.

We devised a new metric called the area ratio that takes the value of 1 for the correlation-based neurons and 0 for the match-based neurons (Figure 4B). The numerator of the area ratio is the area (integral) of the signed amplitude ratio function across the range where the function is negative (inverted tuning); the denominator is the area of the range where the function is positive. Therefore, the area ratio quantifies how strongly each neuron encodes the disparity with the inverted tuning shape as opposed to the non-inverted tuning shape. We used the amplitude parameters of Gabor functions fitted to disparity-tuning data to calculate the signed amplitude ratios. Then, we fitted a quadratic function to interpolate the signed amplitude ratios across binocular correlations. The example MT and V4 neurons shown in Figure 2 had the area ratios of 0.79 and 0.15, respectively (Figures 4C,D).

The distributions of the area ratio markedly differed between MT and V4 (density distributions, Figures 5A,B; cumulative distributions, Figure 5C). The distribution for MT was bimodal: an approximately normal distribution centered at around 0.6 and another peak at 0. The distribution for V4 was unimodal and was strongly skewed towards 0. Quantile–quantile plots revealed that the MT and V4 data were close to normal and exponential distributions, respectively, within the interquartile range (Figure S2). The shape difference was statistically significant (Figure 5C; Kolmogorov-Smirnov test, *p* = 7.9×10^−5^). In MT, the median area ratio was 0.50 and significantly larger than 0 (two-sided Wilcoxson signed-rank test, *p* = 1.6×10^−10^). In V4, it was 0.17 and significantly lower than that in MT (two-sided Mann-Whitney *U*-test, *p* = 3.6×10^−4^). These results suggest that MT and V4 are qualitatively different in terms of the composition of disparity-selective neurons. In area V4, many single neurons are specialized for the match-based representation. In area MT, the distribution seems bimodal: some have the match-based representation but many have representations that are intermediate between fully match-based and fully correlation-based.

Critically, the difference between MT and V4 was revealed only by the area ratio and not by the amplitude ratio, the latter of which is the conventional metric based only on the responses to cRDSs and aRDSs. The distribution of the amplitude ratio was indistinguishable between the two areas (Figures 5D–F). The median values were 0.32 and 0.30 for MT and V4, respectively (two-sided Mann-Whitney *U*-test, *p* = 0.47). The results of this control analysis are instructive, because although there were previous speculations regarding the difference between MT and V4 in disparity representation, these were based on the neuronal response to aRDSs (Krug et al., 2004; Tanabe et al., 2004; for a review, see Parker, 2007). Our results show that characterizing the disparity selectivity at several intermediate levels of binocular correlation between −100% and 100% was necessary to dissociate the depth representations in MT and V4.

The above analyses were performed for the neurons whose responses were well fit by Gabor functions (for disparity-tuning curves) and quadratic functions (for the signed amplitude ratio). We devised similar analyses that did not rely on the model fitting and confirmed the original conclusion (Figure S3A, two-sided Mann-Whitney *U*-test, *p* = 2.3×10^−5^ for the median difference of the model-free area ratio between MT and V4). Moreover, the difference between V4 and MT persisted even when we confined our analyses to the data obtained from the same individual monkey (Figure S3B, two-sided Mann-Whitney *U*-test, *p* = 9.9×10^−3^).

We used dynamic RDSs in which dots randomly appeared and disappeared without any coherent motion, whereas most previous studies on stereo processing in MT used moving dots with 100% motion coherence (e.g., DeAngelis and Uka, 2003; Krug et al., 2004). One might argue that the use of non-coherent-motion RDSs in our experiments resulted in poorer disparity selectivity and led to a biased estimation towards the correlation-based representation in MT. This explanation is possible because stimuli with non-preferred motion parameters decrease the strength of disparity selectivity for cRDSs in MT (Palanca and DeAngelis, 2003). At least in V1, neurons with weaker disparity selectivity for cRDSs exhibit responses closer to the correlation-based prediction (Henriksen et al. 2016b). However, we found no correlation between area ratio and the disparity selectivity for cRDSs (MT: Spearman’s correlation coefficient, *r_s_* = −0.25, p > 0.05; V4: *r_s_* = −0.25, *p* > 0.1; **Figures S4A,B**), or between area ratio and the baseline response (MT: *r_s_* = −0.12, *p* > 0.3; V4: *r_s_* = −0.08, *p* > 0.6; **Figures S4C,D**). These results suggest that neither the strength of the disparity selectivity nor the responsiveness is strongly related to which type of disparity representation a neuron exhibits in MT and V4. Therefore, we believe that the use of non-moving RDSs was unlikely to bias our estimation of area ratios, even though these RDSs might not drive MT neurons to their maximum responsiveness.

### Disparity-tuning symmetry predicts disparity representation in MT and V4

Can we predict if a disparity-selective neuron has a match-based representation or correlation-based representation based on its responses to cRDSs? If so, what response feature carries the information about the representation type? We reported above that the strength of disparity selectivity or responsiveness was not correlated with the area ratio. Here, we examined the shape of the disparity-tuning curve obtained with cRDSs. The disparity-tuning curve has a more complicated form than orientation or direction-tuning curves, with an additional symmetry (phase) parameter. The tuning-curve symmetry has been of particular interest because it reflects the underlying mechanism of disparity detection and implies the utility of the detected signals (Cumming and DeAngelis, 2001; DeAngelis et al., 1991; Marr and Poggio, 1979; Poggio and Fischer, 1977). For example, even-symmetric disparity tuning is a strong indication that the receptive fields in the left and right eyes have the same shape with a position offset (Ohzawa et al., 1997; Tanabe and Cumming, 2008). This mirrors the fact that an object located in depth generates a positional offset when projected to the left and right eyes. Therefore, even-symmetric disparity-selective neurons could be most suited to represent naturally occurring configurations of binocular images. Conversely, odd-symmetric tuning implies that the receptive fields have different shapes (phases) between the two eyes (DeAngelis et al., 1991; see Heafner and Cumming, 2008; Tanabe and Cumming, 2008 for a more complete model). These neurons do not maximally respond to natural binocular images since object’s natural projections to the left and right eyes do not substantially differ in shape. Thus, odd-symmetric neurons could serve as the detector for an impossible pair of binocular images stimulating their receptive fields, thereby providing false-alarm signals to other populations of neurons (Read and Cumming, 2007).

We found a strikingly general relationship between the symmetry of the disparity-tuning curve and the type of disparity representation: neurons with even-symmetric tuning tended to have the match-based representation within both MT and V4. In addition, this relationship applied to the overall comparison between MT and V4. To quantify the symmetry of a tuning curve, we devised a metric that monotonically encodes the tuning-curve symmetry (see reflected symmetry phase in the Methods section; see Read and Cumming 2004 for symmetry phase). A larger value of the reflected symmetry phase indicates a stronger odd symmetry, with 0 for the pure even-symmetric shape (“tuned excitatory” or “tuned inhibitory” types by a conventional classification) and π/2 for the pure odd-symmetric shape (“near” or “far” types). The area ratio was positively correlated with the reflected symmetry phase both in MT and V4 (Figures 6A,B; MT: *r_s_* = 0.55, *p* = 4.0×10^−6^; V4: *r_s_* = 0.34, *p* = 0.025). The analyses based on the fitted Gabor phase instead of the reflected symmetry phase confirmed the same conclusion (**Figures S5A,B**).

**Figure 6.**
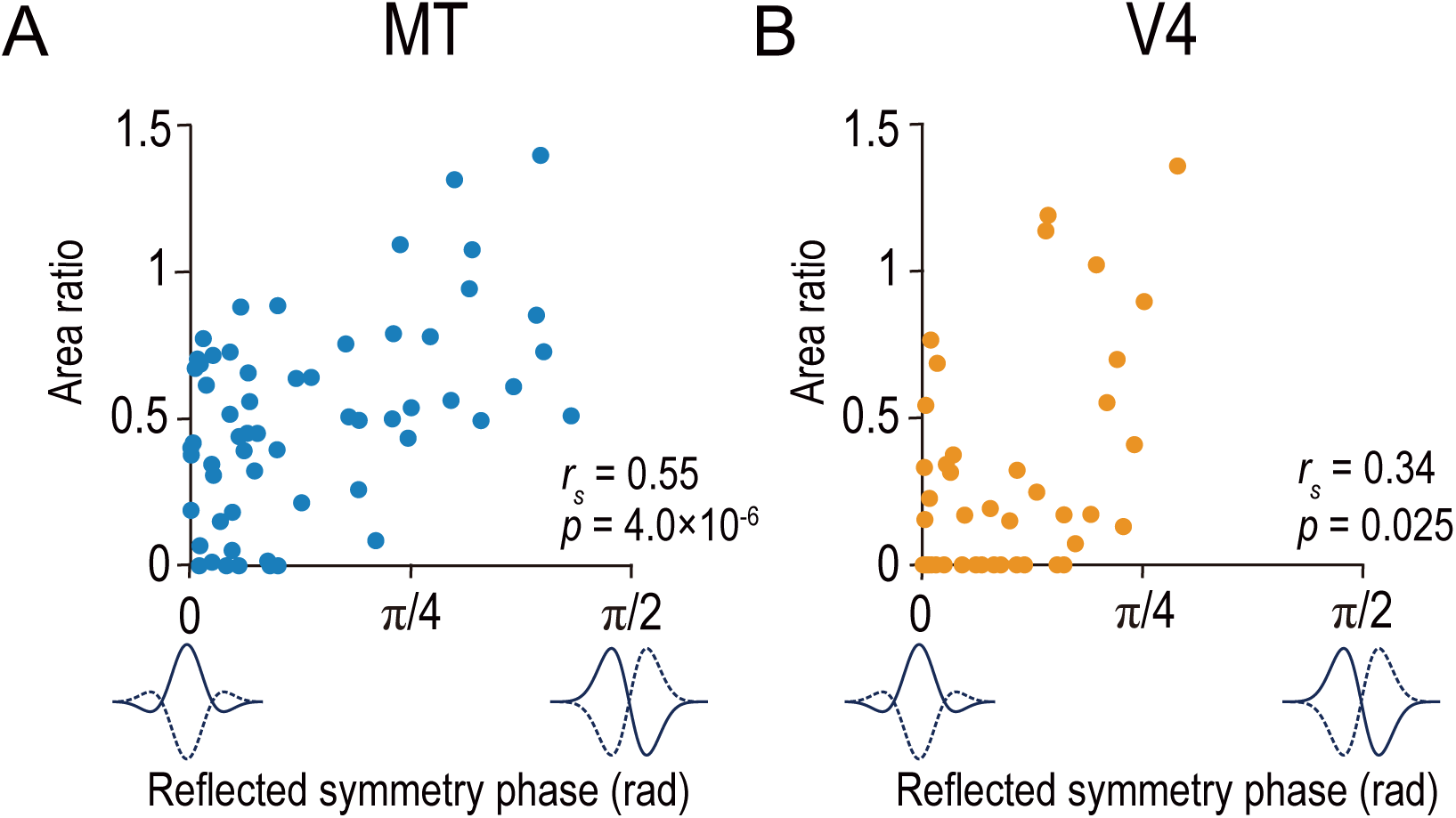
Area ratio is correlated with tuning-curve symmetry in both MT and V4. Area ratios of neurons in MT (**A**) and V4 (**B**) were plotted against the reflected symmetry phase, which takes the value of 0 and π/2 for even-symmetric and odd-symmetric tuning curves, respectively. The symmetry was computed for the responses to correlated RDSs. The plots consist of the neurons with good fitting quality (R^2^ > 0.6 both for Gabor functions for disparity-tuning curves and quadratic functions for the signed amplitude ratio; N = 59 and 43 for MT and V4, respectively). *r_s_* indicates Spearman’s correlation coefficient.

The same relationship also dictated the overall difference between neurons in MT and V4. We showed that neurons in MT had larger area ratios than those in V4 (Figure 5C). We found that neurons in MT had more strongly odd-symmetric tuning curves than those in V4 by comparing their distributions of the reflected symmetry phase (Figure 7). In both MT and V4, the phase was biased toward even-symmetric shapes: the histograms had peaks near 0 (Figures 7A,B). However, the distribution for MT was less strongly biased toward the even shape (a reflected symmetry phase of 0) than that for V4. The difference was clearer in the cumulative distribution (Figure 7C). Although the median values happened to be almost identical (0.074π and 0.073π for MT and V4, respectively), the cumulative distribution for V4 was consistently biased towards smaller values than that for MT. Furthermore, the quantiles of these two distributions had a linear relationship with a slope value of 1.39 (Figure 7D; SE = 0.015; R^2^ = 0.98; MT quantiles plotted against V4 quantiles), meaning that multiplying the reflected symmetry phases for the V4 data set by 1.39 brings the V4 and MT distributions into alignment. Therefore, the disparity tunings in V4 had more even-symmetric shapes than those in MT. At the same time, the disparity representations in V4 were more match-based than those in MT. These results yield a strikingly general principle of the stereoscopic system both within and across the dorsal and ventral pathways: neurons with even-symmetric disparity-tuning curves preferentially constitute the neural solution to the correspondence problem compared to those with odd-tuning curves.

**Figure 7.**
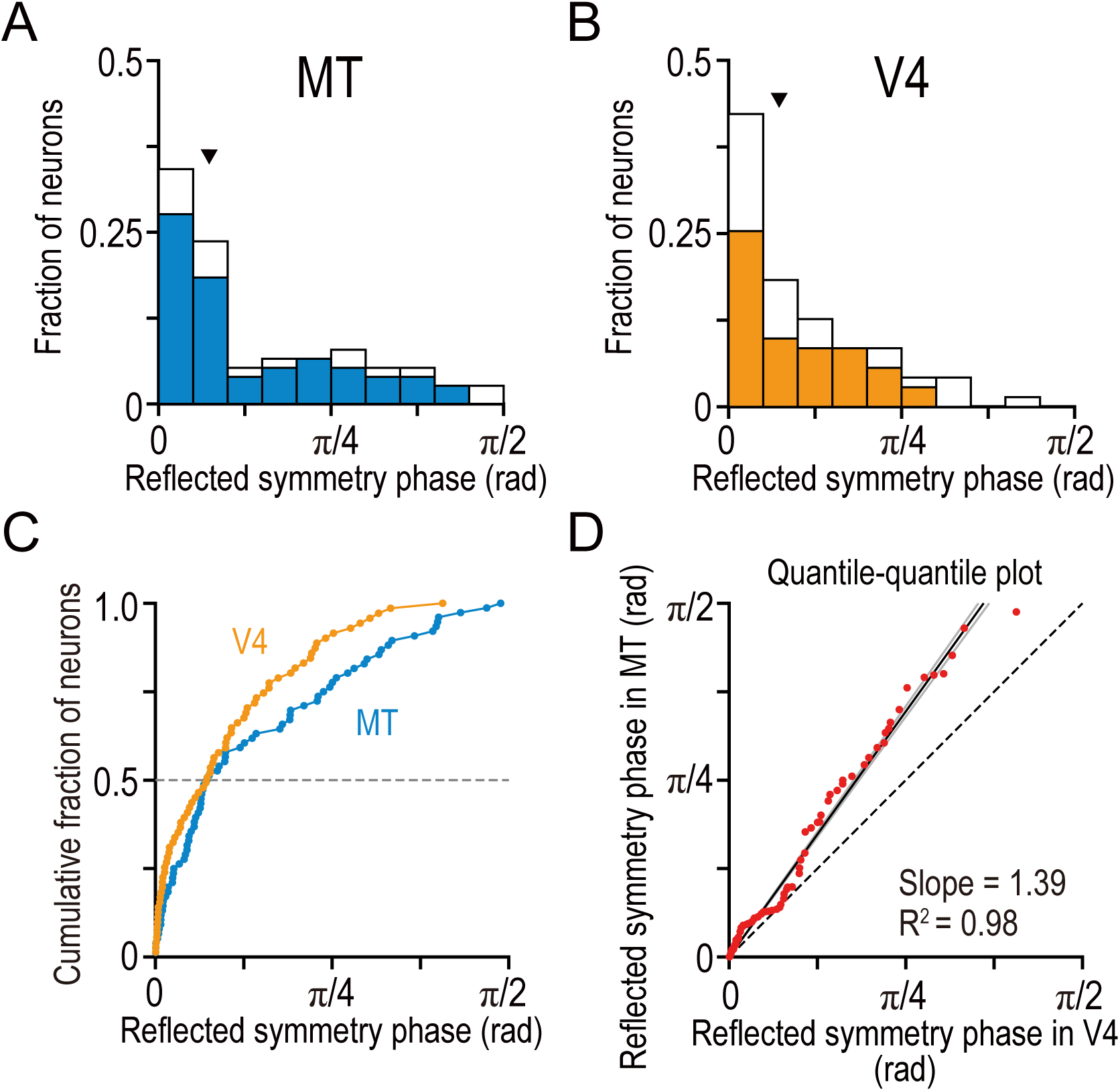
Tuning-curve symmetry is more strongly biased toward the even shape in V4 than in MT. (**A, B**) The distributions of the reflected symmetry phase. We included the neurons for which Gabor functions were well fitted to the disparity tuning (R^2^ > 0.6; N = 76 and 71 for MT and V4, respectively). The parts of the histograms with solid color correspond to the neurons for which quadratic functions were well fitted to the signed amplitude ratio (i.e., the same subpopulations as plotted in Figure 6 and Figures 5A–C). Triangles indicate the medians. (**C**) Cumulative distributions of the reflected symmetry phase for the full populations shown in **A** and **B**. (**D**) Quantile-quantile plot comparing the shape of the distribution between V4 and MT. N = 71 (the number of V4 neurons, which was smaller than that of MT neurons). The horizontal axis plots the sorted values of the reflected symmetry phase for V4. The vertical axis plots the interpolated values of the reflected symmetry phase for MT at the corresponding quantiles. The black line indicates the linear regression. The flanking gray lines indicate the 95% confidence interval of the linear regression.

### MT exhibits stronger and faster disparity selectivity than V4 for correlated RDSs

What is the advantage of the disparity signals transmitted by MT? We showed that MT had a more primitive, correlation-based disparity representation than V4, which manifested in how the disparity selectivity changed with graded anticorrelation. Here we show that the disparity selectivity for cRDSs is faster and stronger in MT than in V4 (Figure 8). We devised a metric called signed disparity discrimination index (sDDI) and computed it using a sliding time window both before and during the stimulus presentation (10-ms width and step; see Prince et al., 2002 for DDI). The absolute value of sDDI indicates the strength of the selectivity relative to the response noise: 0 for no selectivity and 1 for extremely large selectivity compared to the noise level. The positive and negative signs indicate whether the instantaneous selectivity quantified over a short time window is consistent with or opposite from the overall selectivity quantified over the whole stimulus duration, respectively.

**Figure 8.**
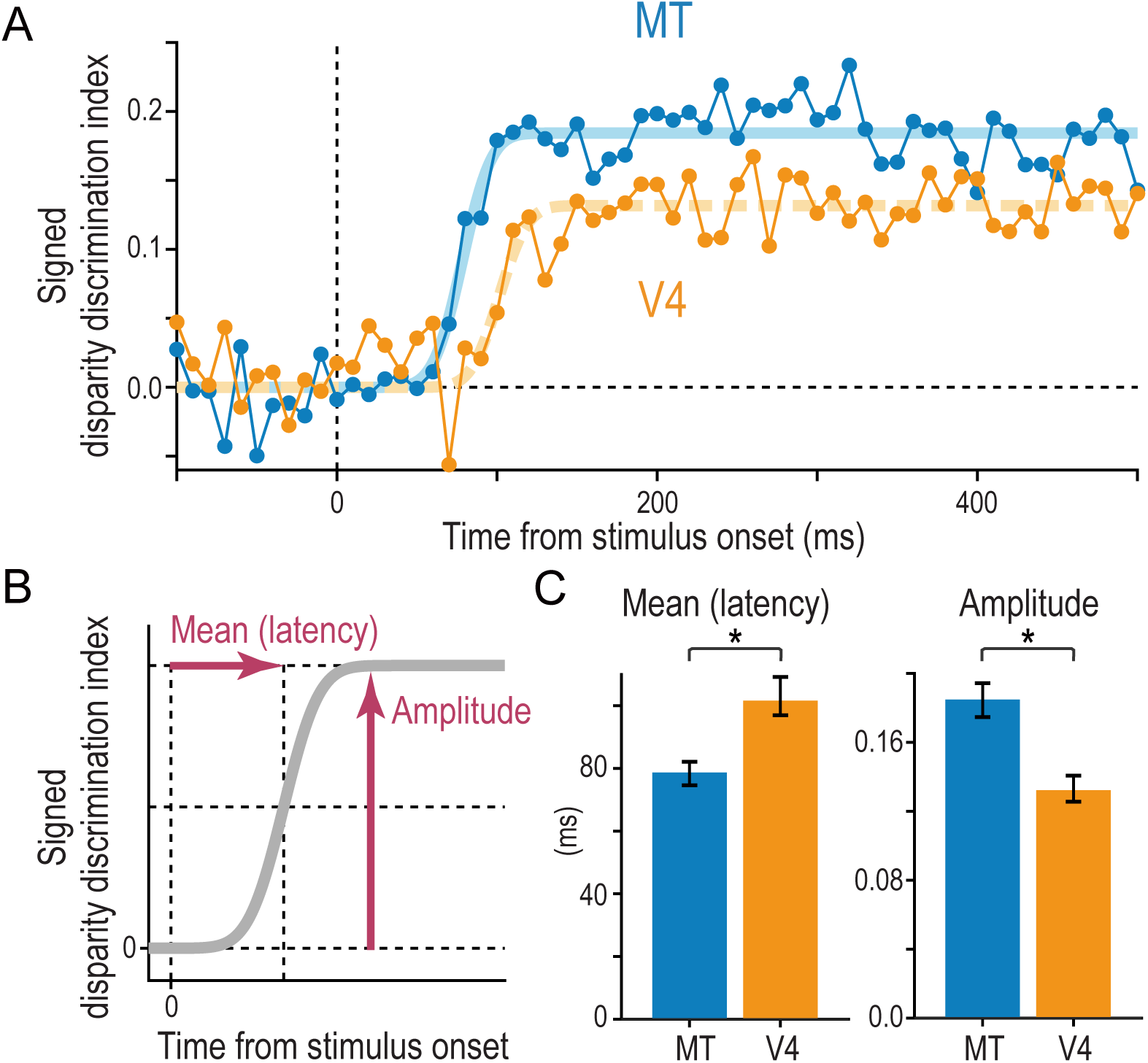
Comparison of the strength and speed of disparity selectivity between MT and V4 in response to cRDSs. (**A**) The average time course of the signed disparity discrimination index (sDDI). The sDDI was calculated within 10-ms (non-overlapping) time windows for each cell and then averaged across cells separately for MT and V4 (N = 83 and 78, respectively). The smooth curves are fitted cumulative Gaussian functions. The vertical dashed line indicates the stimulus onset. (**B**) Parameters of the fitted cumulative Gaussian function. The amplitude parameter quantifies the saturation level of the sDDI. The mean parameter, our estimate of the selectivity latency, corresponds to the time required for the sDDI to rise to half the final amplitude. (**C**) Best-fitted values and 68% bootstrap confidence intervals. The latency and amplitude were significantly different between MT and V4.

We found that the average sDDI of MT neurons rose faster and reached a higher level than that of V4 neurons (Figure 8A). We fitted the cumulative Gaussian function to estimate the latency and amplitude of the disparity selectivity (mean and amplitude parameters, respectively; Figure 8B). The latency was 78 ms in MT and 101 ms in V4 (Figure 8C). The peak sDDI was 0.18 in MT and 0.13 in V4. Thus, compared with MT, in V4 the latency was 29% greater and the strength was 28% lower. The differences were statistically significant (*p* = 0.0083 and *p* < 1.0 × 10^-4^ for the latency and amplitude differences between MT and V4, respectively; 10,000 bootstrap resamples). These results were surprising given that we used the RDSs suitable for driving V4 neurons (Abdolrahmani et al. 2016; Kumano et al., 2008; Shiozaki et al., 2012; Tanabe et al., 2005; Tanabe et al. 2004; Umeda et al. 2007). For example, unlike previous MT studies (DeAngelis and Uka, 2003; Krug et al. 2004), our RDSs did not contain visual motion, and their dot patterns were updated only slowly (10.6 Hz). Overall, we suggest that MT signals disparity more robustly and faster than V4, providing signals useful for rapid three-dimensional eye movements (Masson et al., 1997) and urgent depth judgments in response to correlated stimuli.

## Discussion

We used graded anticorrelation to identify whether MT and V4 neurons represented disparity based on binocular correlation (as prescribed by the disparity energy model; Ohzawa et al., 1990) or on binocular match (as consistent with the solution to the binocular correspondence problem) (Figure 1). An area-wise characterization of disparity tuning detected the signatures of the correlation-based representation in MT and those of the match-based representation in V4 (Figure 3). A finer, cell-by-cell characterization revealed a qualitative difference in the distribution of the representation type between the two areas (Figures 5A–C). MT appears to consist of two groups of neurons: one with the match-based representation and the other with the intermediate representation (Figure 5A). V4 lacked the intermediate group (Figure 5B). This dissociation disappeared with the analysis based on the conventional methods (Figures 5D–F). Importantly, in both areas, the shape of the disparity tuning (in responses to cRDSs) predicted the type of representation: more even-symmetric neurons showed stronger biases toward the match-based representation (Figure 6). This relationship was also observed for the inter-area comparison between MT and V4 (Figures 5,7). Lastly, although the type of disparity representation was more primitive overall in MT than in V4, the disparity signaling for cRDSs was stronger and faster in MT (Figure 8).

### Explanations for the links between even-symmetric disparity tuning and match-based disparity representation

The symmetry (phase) of disparity-tuning curves has been studied for decades to understand the neural coding of disparity (DeAngelis et al., 1991; DeAngelis and Uka, 2003; Hinkle and Connor, 2005; Ohzawa et al., 1997; Poggio and Fischer, 1977; Poggio et al., 1988; Prince et al., 2002; Read and Cumming, 2004; Tanabe and Cumming, 2008; Tanabe et al., 2005). Although many theories have been proposed (e.g., Fleet et al., 1996; Goncalves and Welchman, 2017; Read and Cumming 2007), no empirical data have been reported regarding how the disparity-tuning symmetry relates to the process of solving a fundamental computational problem of stereopsis—the correspondence problem. In this study, we found that neurons with even-symmetric tuning preferentially constituted the neural solution to the correspondence problem in extrastriate areas (Figure 6). Below we propose two interpretations regarding this novel finding.

First, we propose that the observed link between tuning symmetry and disparity representation may share a common cause: adaptation of the visual system to the natural configurations of binocular images. The correlation-based representation is the product of a simple mechanism that does not consider whether the input images have natural or unnatural configurations. By contrast, the match-based representation can be viewed as the result of the active adaptation to naturally occurring binocular images (Haefner and Cumming, 2008). Within our stimulus space, the strength of the binocular match indicates how naturalistic our stimuli are: cRDSs are most naturalistic in the sense that the correspondence problem has a smooth solution (Marr and Poggio, 1976); aRDSs are naturally impossible stimuli, and half-matched RDSs are an intermediate type (Figure 1). In parallel to the type of disparity representation, the symmetry of disparity tuning also carries implications regarding the process of natural versus unnatural binocular images. Neurons with even-symmetric disparity tuning are better suited to detect binocular images with natural configurations. This is because the left-eye and right-eye receptive fields of these neurons have a positional offset, just as natural pairs of binocular images do. By contrast, neurons with odd-symmetric tuning are bound to respond most vigorously to unnatural binocular images (Read and Cumming, 2007), because the monocular receptive fields of odd neurons have interocular phase differences (for energy model-based analyses, see DeAngelis et al., 1991; Ohzawa et al., 1997; Prince, Cumming, and Parker, 2002; for experimental verification in macaque V1 and V2, see Tanabe and Cumming, 2008). Such interocular phase shifts do not occur in natural binocular stimuli.

We have explained the adaptation to natural binocular inputs in two different aspects of neuronal disparity selectivity: disparity representation and disparity-tuning symmetry. These two aspects could be independent characteristics of cortical disparity coding. In fact, according to the disparity energy model, all disparity detectors should have the correlation-based representation irrespective of the tuning symmetry. Nevertheless, we repeatedly found the same link between disparity representation and tuning symmetry within V4, within MT, and across the two areas. A network of extrastriate areas may converge the two kinds of adaptation to natural binocular inputs onto the same subpopulation of disparity-selective neurons. Such neurons can be viewed as having an efficient representation of binocular disparity because they dedicate their disparity encoding to the naturally occurring inputs (Attneave, 1954; Barlow, 1961; Haefner and Cumming, 2008).

The second interpretation regards the mechanisms for transforming the simple correlation-based representation into the match-based representation. Adding expansive nonlinearity to disparity energy models can achieve the transformation under some conditions (Doi and Fujita, 2014; Lippert and Wagner, 2001). Although the additional nonlinearity must be only part of the full mechanism, it has psychophysical (Doi et al., 2013; Henriksen et al., 2016a) and physiological support (Henriksen et al., 2016b). A key constraint of this mechanism is that the energy-model units should have even-symmetric, but not odd-symmetric, disparity tuning. For example, the tuning shape for hmRDSs is consistent with that for cRDSs only when the initial units have even-symmetric tuning (Doi et al., 2013). This constraint could also explain why the match-based representation is more prevalent in neurons with even-symmetric tuning.

### Contributions of the MT and V4 disparity signals to perception, eye movements, and downstream activity

Two different laboratories have independently examined MT and V4 with aRDSs before. A naive comparison of their results suggests that neurons in V4 have response patterns closer to the correspondence solution than those in MT (compare Figure 3C in Krug et al., 2004 and Figure 6A in Tanabe et al., 2004; for review see Parker, 2007). However, the apparent difference between MT and V4 in the two studies might arise from differences in the cell-selection procedures used: the MT study excluded the cells without the significant disparity selectivity for aRDSs from the figure, whereas the V4 study did not. Indeed, a more recent study showed that when the selection procedure was matched, the population-averaged response to aRDS was nearly indistinguishable between MT and V4 (Abdolrahmani et al., 2016). We confirmed this conclusion in the present study, where the experimental and analysis methods were matched between the two areas more than any previous comparison (Figures 5D–F).

Using a novel analysis method, we found a qualitative difference in disparity representations between MT and V4 (Figures 5A–C). What are the roles of each area in perception and action? We propose that MT pathway quickly and robustly relays the correlation-based disparity signals to the behavioral stage. The use of MT’s disparity signals should not be limited to visually guided actions, because the link between the disparity-selective neurons in MT and stereoscopic depth judgment has been demonstrated extensively using a coarse depth task (Chowdhury and DeAngelis, 2008; DeAngelis et al. 1998; Uka and DeAngelis, 2004).

Interestingly, MT neurons with more odd-symmetric tuning have stronger response correlation (i.e., choice probability) to the judgment of coarse depth (Uka and DeAngelis, 2004). This may appear contradictory to our finding that more even-symmetric neurons tended to have the match-based disparity representation. We present two explanations for this apparent contradiction. First, extensive behavioral training probably caused the high choice correlation of odd-symmetric neurons observed in Uka and DeAngelis (2004) (as suggested by the authors). During the training, they used near and far stimuli that are most suitably processed by odd-symmetric neurons. By contrast, our monkeys were trained only on the fixation task, so that the link between the tuning symmetry and representation type reported in this study is an innate property of the stereoscopic system. Second, some aspects of stereoscopic depth perception may not require the correspondence problem to be fully solved; these include coarse stereopsis (considering Uka and DeAngelis [2004] and this study together) and the perception of 3D structure-from-motion (Krug et al., 2004). According to this notion, the correlation-based disparity signals in MT should be able to contribute to depth perception.

Our above proposal is supported by earlier psychophysical studies with the same graded anticorrelation paradigm (see Fujita and Doi, 2016 for a review). Human depth judgment does indeed exhibit a trace of the correlation-based disparity representation (e.g., reversed depth perception for aRDSs; Aoki et al., 2017; Tanabe et al., 2008). Moreover, the inferred contribution of the correlation-based representation to perceived depth increases with the magnitude of stimulus disparity (Doi et al., 2011) as well as with the stimulus pattern refresh rate (Doi et al., 2013). The magnitude effect is consistent with the finding that MT contributes to coarse, but not fine, depth judgment (Uka and DeAngelis, 2006). The refresh-rate effect is consistent with the notion that MT is likely to contribute to perception more with dynamic stimuli than with static ones.

We also suggest that the disparity signals in MT drive the reflexive vergence eye movement via the medial superior temporal area (Takemura et al., 2001). For anticorrelation, the vergence eye movement responds in the opposite direction from stimulus disparity with a reduced amplitude (Masson et al., 1997), suggesting that the driving signal is intermediate between the match-based and correlation-based representations. Many MT neurons we observed fell into this category (see the second peak in the histogram in Figure 5A). The vergence response can be very fast, starting to rise only 60–70 ms after the stimulus onset (Masson et al., 1997), and requiring the fast and robust disparity signals that we observed in MT (Figure 8). Our suggestion is also consistent with the finding that MT neurons represent the “absolute” disparity of a stimulus, without being sensitive to the disparity of a reference plane (Uka and DeAngelis, 2006). This response property is a prerequisite for the vergence drive (Westheimer and Mitchell, 1956).

We suggest that V4 plays a key role in transforming the disparity signals from early visual areas into the inputs useful to the inferior temporal (IT) cortex, thereby helping create the full-fledged representations of 3D objects and 3D scenes at the end stage of the ventral pathway (Yamane et al., 2008; Verhoef et al., 2016). Consistent with the idea that V4 is in the middle of the transformation process is the finding that the correspondence problem is not yet fully solved in the single-neuron activity we observed in V4. Close inspection of Figure 3B reveals that the fraction of disparity-selective neurons did not noticeably decrease from correlation levels 0% to −100%, although the strength of the binocular match decreased from 50% to 0% (Figure 1). This discrepancy can be mitigated by aggregating the population responses just the way decision-making circuits would do (Abdolrahmani et al., 2016). This is because at −100% correlation, roughly half of the V4 neurons had inverted tuning curves (Figure 3D). If the responses of V4 neurons are aggregated according to their preferred disparity for cRDSs (in our stimulus space, these are the most naturalistic stimuli to which the decision mechanisms are likely to adapt), their tuning curves at −100% correlation are averaged out and flattened (Doi et al., 2018). This averaging out would not occur at 0% correlation, because the tuning shape at 0% correlation tended to be consistent with that at 100% correlation (as predicted by models of the match-based representation; Doi and Fujita, 2014; Doi et al., 2013). Thus, the aggregated V4 responses should resemble the pure match-based representation of disparity. This property of V4 neurons should presumably help downstream areas such as the IT cortex represent a complete solution to the correspondence problem at the individual-neuron level (Janssen et al. 2003).

We also suggest that V4 is critical for the perception of fine depth, because fine disparity perception entirely relies on the match-based representation (Doi et al., 2011). Several physiological results strongly support this view. First, V4 neurons have choice-predictive responses during a fine depth task (Shiozaki et al., 2012). Second, microstimulation of a cluster of V4 neurons biases the judgment of fine depth in the way predicted from the response property of the stimulated cluster (Shiozaki et al., 2012). Lastly, V4 neurons encode the “relative” disparity between the center and surround of their receptive fields, a property necessary for supporting stereoacuity and a precursor for fixation-depth-invariant representation of 3D objects (Umeda et al., 2007). We also suggest that V4 is important for 3D shape perception, because the perceptual discrimination of disparity-defined 3D shapes is difficult without match-based disparity signals (Asher and Hibbard, 2018; Tanabe et al., 2008). This is broadly consistent with the finding that V4 is involved in the encoding of 3D shape cues such as 3D slant (Hinkle and Conner, 2002). Overall, the signals in V4, including the selectivity for match-based disparity, fine disparity, and 3D shape cues, should help the ventral pathway derive a fully developed representation of 3D objects and 3D scenes in its highest stage, the IT cortex.

### Concluding remarks

We report the division of labor between extrastriate areas MT and V4 in stereopsis, as revealed by a novel manipulation of RDSs. We suggest that area MT, a mid-tier stage in the dorsal pathway, relays the primitive correlation-based disparity signals without considerably refining the nature of the signals. Instead, this pathway has the advantage in terms of signaling speed and strength, which are useful for rapid vergence eye movements and coarse depth perception. By contrast, area V4, a counterpart stage in the ventral pathway, predominantly processes more sophisticated disparity signals based on binocular match. These signals help the end stage of the ventral pathway, the IT cortex, to fully solve the binocular correspondence problem and derive the fine representations of 3D objects. Therefore, the division of labor between the dorsal and ventral visual pathways may embody the tradeoff between the rapid and robust transmission of sensory signals and the complex computation needed to derive elaborate sensory representations. This proposal provides a unified account for two earlier theories on the dorsal versus ventral division of labor (Goodale and Milner, 1992; Ungerleider and Mishkin, 1982). For example, vision related to action may depend more on the dorsal pathway because it benefits from rapid, robust sensory transmission, and so does visual motion perception. The analyses of colors and objects depend more on the ventral pathway because they require complex computations.

We also note that the response characteristics of individual neurons are diversely distributed across MT and V4. Moreover, MT and V4 share a common link between the disparity-tuning shape and the type of disparity representation. It is possible that the interactions between the dorsal and ventral pathways play several roles in solving the correspondence problem, deriving 3D object representation, making decision about stereoscopic depth, and generating object-oriented motor action (e.g., reaching and grasping), as suggested by a broad range of evidence from neuroanatomy, single-neuron recording, electro-encephalography, psychophysics, and brain imaging (Borra et al., 2008, 2010; Cottereau et al., 2014; Farivar, 2009; Fujita and Doi, 2016; Freud et al., 2016; Janssen et al., 2018; Van Polanen and Davare, 2015).

## Acknowledgments

We thank H. Ban, Y. Sakano, H.M. Shiozaki, and B.G. Wundari for comments on earlier versions of the manuscript; S. Mita, T. Oga, M. Onoue, and M. Inagaki for technical assistance; T. Uka, K. Okada, Y. Kobayashi, H. Kumano, T.M. Sanada, and M. Saruwatari for technical advice; and M. Omokawa for animal care. This work was supported by grants to I.F. from the Ministry of Education, Culture, Sports, Science and Technology of Japan (MEXT; 23240047, 15H01437, 17H01381, 18H05007) and the Ministry of Internal Affairs and Communications. The monkeys were provided by the National Institute of Natural Sciences (NINS) through the National Bio-Resource Project (NBRP) of the MEXT, Japan.

## Author Contributions

T.W.Y., T.D., M.A., and I.F. designed the research; T.W.Y. and M.A. conducted the MT and V4 experiments, respectively; T.W.Y. and T.D. analyzed the data; and T.W.Y., T.D., M.A., and I.F. wrote the paper.

## Competing Interests

The authors declare no competing interests.

## Methods

We recorded single neuron responses from areas MT and V4 of two Japanese monkeys (*Macaca fuscata*). All procedures were approved by the Animal Experiment Committee of Osaka University, and conformed to the Guide for the Care and Use of Laboratory Animals issued by the National Institutes of Health, USA. The data sets and analysis codes for reproducing the results reported in this study are available upon reasonable request to the lead contact Ichiro Fujita.

### Animal preparation

We used two Japanese monkeys (Monkey O, male, weighing 8.5 kg; Monkey A, female, weighing 5.7 kg). In both monkeys, a head-holding device was attached to the skull and Teflon-insulated stainless-steel search coils were implanted between the conjunctiva and sclera in both eyes under anesthetic and aseptic conditions. For details of the surgical procedures, see Kumano et al. (2008). Briefly, the monkeys were administered atropine sulfate (0.02 mg/kg, i.m.) to reduce salivation and were sedated with ketamine (5 mg/kg, i.m.). Anesthesia was maintained by inhalation of a mixture of isoflurane (0.3–2.0%, Forane, Abbott), nitrous oxide (66%), and oxygen (33%). Lidocaine was used for local anesthesia as needed. Throughout surgery, we continuously monitored electrocardiogram data, arterial oxygen saturation levels (SPO2), end-tidal CO_2_ levels, and heart rate. Postoperatively, we administered a general antibiotic (piperacillin sodium), ocular antibiotic (ofloxacin), glucocorticoid (betamethasone), and analgesic (ketoprofen). We trained the monkeys to perform a fixation task (see below). When the training was complete, we performed another surgery under the same anesthetic procedure to implant a chamber for recording from MT (inner diameter, 19 mm; Crist Instruments, Hagerstown, MD). The chamber was placed on the occipital cortex 17 mm lateral and 14 mm dorsal to the occipital ridge tilted posteriorly at an angle of 25° above the horizontal. The skull inside the chamber was removed with a trephine (Fine Science Tools, North Vancouver, Canada). We collected the data of V4 neurons in a previous study (Abdolrahmani et al., 2016) by placing a recording chamber 25 mm dorsal and 5 mm posterior to the external ear canals. For the V4 experiments, we used Monkey O (the same monkey used in MT recording) and Monkey I (*Macaca mulatta*, male) weighing 6.5 kg.

### Behavioral task

The monkeys viewed a CRT monitor (refresh rate 85 Hz; Multiscan G520, Sony) at a distance of 57 cm. The monitor covered a visual angle of 40° × 30°. We alternately presented images to the left and right eyes with custom-made OpenGL software, a quad-buffered graphics card, and shutter glasses based on ferroelectric liquid crystal devices (LV2500P-OEM, Citizen FineDevice, Yamanashi, Japan). The left and right shutter glasses alternately closed at each refresh of the display. Although the use of the shutter glasses decreases the effective binocular correlation compared with a haploscope, we have previously shown that the effective correlation is an unbiased, linear function of the notional correlation value (Abdolrahmani et al., 2016). Therefore, the use of shutter glasses did not complicate our interpretations as a result of artifactual distortion of the effective binocular correlation. Upon presentation of a white point at the center of the monitor, the monkeys were required to fixate it within 500 ms. After fixation, a visual stimulus was presented for 500 ms to both monkeys in the MT experiment and to monkey O in the V4 experiment; the stimulus duration was 700 ms for monkey I in the V4 experiment. The monkeys had to maintain fixation within a 1.5° × 1.5° window and a vergence angle within ± 0.25° for another 200 ms. The eye positions were monitored with a scleral eye-coil system (DSC-2000, Sankei Kizai, Tokyo, Japan). For successful trials, the monkeys received drops of juice or water as a reward. When the monkeys failed to maintain fixation, they did not receive a reward, and the data were discarded.

### Visual stimuli

We used circular patches of dynamic random-dot stereograms (RDSs) as visual stimuli. The RDSs consisted of a central disk and a surrounding ring with no gap between them. The central disk was positioned to cover the classical receptive field (RF) of a recorded neuron. The surrounding ring was 1.6° wide. RDSs contained an equal number of bright (2.28 cd/m^2^) and dark (0.01 cd/m^2^) dots with a gray background (1.14 cd/m^2^). Luminance was measured through the shutter glasses placed between the monitor and a photometer (CS1000, Konica Minolta, Tokyo, Japan), and linearized (gamma corrected). We used only red phosphors to minimize inter-ocular crosstalk (< 3% of the background) because the decay time is shorter for red phosphors than for green and blue phosphors. The dot size was 0.17° × 0.17° with anti-aliasing. The dot density was 25%.

We manipulated the binocular disparity and binocular correlation of the central disk, whereas these aspects of the surrounding ring were fixed at zero disparity and 100% correlation. We manipulated the binocular correlation of the central disk by reversing the luminance contrast of a varying proportion of dots in one eye (graded anticorrelation, Figure 1A; Doi et al., 2011, 2013). Note that this manipulation of the stimulus correlation is detectable only through binocular viewing; it does not change the overall luminance of the stimulus, because the RDSs are composed of the same numbers of bright and dark dots.

### Recording techniques

A tungsten electrode (0.3–2.0 MΩ at 1 kHz; FHC Bowdoinham, ME) was inserted into the cortex through a transdural guide tube using a micromanipulator (MO-971, Narishige, Tokyo, Japan). Extracellular neural signals were amplified (×5,000; MDA4-I, Bak Electronics, Mount Airy, MD) and filtered (0.5–5 kHz; Multifunction Filter 3624, NF Corporation, Yokohama, Japan). Action potentials from single neurons were isolated by either a dual-window discriminator (DDIS-1, Bak Electronics, Mount Airy, MD) or a template-matching sorting system (Multi Spike Detector, Alpha-Omega Engineering, Nazareth, Israel). Times of spike occurrences and positions of both eyes were stored at 1-ms resolution for offline analysis.

We identified area MT based on the motion direction tuning of single- and multi-neuron activity, the relation between eccentricity and RF size, and changes in RF location along the electrode penetrations (Gattas & Gross, 1981; Van Essen et al., 1981), as well as general anatomical position. We confirmed the location of MT based on the pattern of gray matter and white matter encountered during electrode penetration before reaching MT as well as during the subsequent entry, after a short silent region (sulcus), into area MST, which had large, contralateral RFs covering a quadrant or half of the display (Gu et al., 2006; Uka & DeAngelis, 2003). After all recording experiments were finished, we made electrolytic lesions (10 nA for 10 s, electrode negative) at three to five sites in or around the recording regions. The monkeys were then euthanized with an overdose infusion of pentobarbital sodium and perfused from the heart with 0.1% phosphate-buffered saline (PBS) and 4% paraformaldehyde. The brains were removed, postfixed, frozen, and cut into 60-micron-thick serial, parasagittal sections. Alternate sections were stained for cell bodies with the Nissl method or for myelinated axons with the Gallyas method (Gallyas, 1979). We successfully recovered all the lesions. We verified that we recorded from MT deep in the posterior bank of a caudal part of the superior temporal sulcus where layers IV–VI contained a dense and uniform band of myelin fibers (Ungerleider and Mishkin, 1979; Van Essen et al., 1981). For the identification and RF information of V4 neurons, see Abdolrahmani et al. (2016).

Most of our MT neurons (65 out of 83) had their RFs in the lower visual field; RFs of all of our V4 neurons (N = 78) were located in lower visual field. The mean eccentricity was 6.9° (SD = 3.3°) for MT and 9.0° (SD = 3.0°) for V4 in our data sets. Although the mean eccentricity was slightly larger in V4 than in MT, we found no correlation between the area ratio (our metric for disparity representation: see below) and eccentricity either in MT (Spearman’s correlation *r_s_* = −0.035, *p* = 0.79) or in V4 (*r_s_* = −0.065, *p* = 0.68). Therefore, it is unlikely that the observed difference in disparity representation between MT and V4 reflects the potentially confounding effects of eccentricity.

### Experimental procedures

Here we describe the procedures used for area MT. We used similar procedures for area V4, but see Abdolrahmani et al., 2016 for details. After isolating single-neuron activity, we determined its classical RF by a patch of RDSs (size: 3° or smaller, disparity: 0°, dot-pattern refresh rate: 10.6 Hz in most cases). We measured direction tuning with a patch of dots coherently moving in one of eight directions (45° apart). We then probed the range of disparity tuning with dynamic, non-motion-coherent RDSs in several (at least three) stimulus presentation trials. For this initial test, we used cRDSs with nine disparities (−1.6° to 1.6° in 0.4° intervals) and uncorrelated RDSs (uRDSs; RDSs with independently created left-eye and right-eye images). After all these tests, we measured the responses to RDSs with several binocular disparities (tailored for each recorded neuron based on the initial test) and correlation levels (100%, 70%, 30%, 0%, −30%, −70%, and −100%). The dot-pattern refresh rate was set to 10.6 Hz, the value used for the V4 experiment.

### Data analysis

All analyses were carried out with MATLAB (The MathWorks). The spontaneous firing rate was calculated for a 250-ms period preceding the stimulus onset and averaged across trials. To construct disparity-tuning curves, the mean firing rate at each correlation level and disparity was calculated for a certain time window. The time window overlapped with the stimulus presentation period but was delayed to take into account response onset and offset latency (60 ms for MT and 80 ms for V4). The average firing rate was based on at least five trials. One of the monkeys (Monkey O) was used in both MT and V4 experiments. The V4 data were used in our previous publication (Abdolrahmani et al., 2016), but the main analyses of the current paper are all new. Particularly, we quantified the disparity representation of each V4 neuron and directly compared it against MT results in this study.

#### Gabor function for disparity tuning

We characterized neuronal disparity tuning by fitting a Gabor function, a product of a Gaussian envelope and a cosine carrier, to the mean firing rate data. At each level of binocular correlation, the mean firing rate (*R*) at disparity (*x*) was defined as follows:

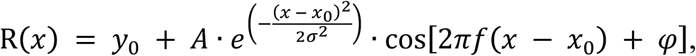

where *A*, *y_0_*, *x_0_*, *σ*, *f* and *φ* represent the amplitude, baseline response, horizontal position, envelope width, carrier frequency, and carrier phase, respectively. We performed a half-wave rectification for *R* because the firing rate cannot be negative. Only the amplitude and phase parameters were independently fitted across different binocular correlation levels, because the differences in these two parameters characterize the correlation-based and match-based representations (Figure 1B). The other parameters were shared across different correlation levels. The fmincon function in the MATLAB Optimization Toolbox was used for finding the parameter combination that minimizes the summed squared error between the single-trial firing rates and Gabor functions. The goodness of fit was estimated by taking the R^2^ between the mean firing rates and the best-fitted Gabor function separately for each correlation level. After fitting, we selected 76 MT and 71 V4 disparity-selective neurons that had good fitting quality for the responses to cRDSs (R^2^ > 0.6).

#### Signed amplitude ratio

To characterize how a binocular-correlation decrease from 100% changes disparity tuning, we devised the signed amplitude ratio by modifying a more conventional metric, the amplitude ratio (Cumming & Parker 1997). We computed the signed amplitude ratio at each level of binocular correlation for each cell. The conventional amplitude ratio quantifies how much the fitted Gabor amplitude changes because of the anticorrelation of stereograms. The conventional metric was given a sign that was determined based on how anticorrelation changed the shape of disparity tuning. When anticorrelation inverted the shape, we gave a negative sign. We detected the shape inversion based on the change in the symmetry phase (see below). The symmetry phase difference between two tuning curves had to be smaller than −0.5π or larger than 0.5π for the shape inversion to be detected.

#### Reflected symmetry phase

When the cycle of the carrier sinusoid is much larger than the Gaussian envelope width, the Gabor phase may not be a good indicator for the true shape of disparity-tuning curves (Read and Cumming, 2004). In this case, the sinusoidal carrier does not constrain the shape of the fitted Gabor function. Accordingly, the neurons with even-symmetric tuning curves can be classified as odd symmetric if we use the fitted value of the carrier phase (confirmed in our data set). We therefore characterized the shape of the disparity-tuning curve by calculating the symmetry phase (Read and Cumming, 2004). This metric directly estimates the relative weights of even-symmetric and odd-symmetric components in the overall shape of the fitted tuning curve. The symmetry phase of zero or π corresponds to the pure even symmetry, whereas the symmetry phase of ± π/2 corresponds to pure odd symmetry. To construct a metric that monotonically quantifies the tuning symmetry, we devised reflected symmetry phase. We changed the symmetry phase to the absolute value. Then, we wrapped the absolute value into the range from zero (purely even symmetric) to π/2 (purely odd symmetric) by reflecting it at π/2.

#### Area ratio

We characterized how well each of the recorded disparity-selective neurons followed the correlation-based or match-based prediction using their responses over the entire range of binocular correlation. This analysis consisted of two steps. First, we characterized how the signed amplitude ratio changed as a function of binocular correlation. Although the simplest examples shown in Figure 4A are linear functions, we fitted a quadratic function to the observed data and this often showed nonlinear trends (Figures 4C,D). The quadratic coefficient was almost always positive, meaning that the nonlinearity was expansive (84% for MT, 90% for V4). We quantified the degree of nonlinearity using the ratio of the quadratic coefficient to the linear coefficient from the best-fitted quadratic function (Britten et al., 1993). The ratio was significantly higher than zero for both MT and V4 (MT: median, 0.02; lower-quartile, 0.0057; upper-quartile, 0.0597; two-sided Wilcoxson signed-rank test, *p* = 3.6×10^−14^; V4: median, 0.02; lower-quartile, 0.0116; upper-quartile, 0.028; *p* = 2.4×10^−13^), validating our use of quadratic functions. The function was constrained to take the value of one at 100% correlation. The sum-squared error was minimized. Second, we calculated the area ratio to quantify how well the fitted quadratic function followed the correlation-based prediction versus the match-based prediction (Figures 4A,B). The denominator of the area ratio was the area (integral) of the fitted quadratic function across the range where the fitted function was positive (gray triangle in Figure 4B). The numerator was the area of the function over the range where the function was negative (blue triangle). Because the correlation-based representation should have perfectly inverted disparity-tuning curves for two binocular correlation values with the same magnitude but opposite signs, it should lead to an area ratio of one. By contrast, the match-based representation should have a flat tuning curve at −100% correlation, leading to an area ratio of zero.

#### Signed disparity discrimination index (sDDI)

The conventional disparity discrimination index (DDI) measures the disparity discriminability of a neuron as follows (Prince et al., 2002):

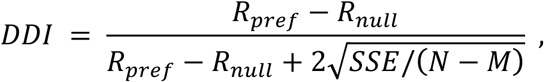

where *R_pref_* and *R_null_* indicate the mean firing rates of a neuron at its preferred and null disparities, respectively (in response to cRDSs). The SSE is the sum of the squared error of single-trial responses (around the mean; i.e., the tuning curve); *N* and *M* indicate the total number of trials and the number of tested disparities, respectively. A DDI of 0 indicates no selectivity (i.e., a completely flat tuning curve); a DDI of 1 indicates that the tuning curve amplitude is so large that the noise level is negligible.

We wanted to compute DDIs using small, sliding time windows to examine how the disparity selectivity developed in MT and V4. Therefore, we modified the conventional DDI as follows:

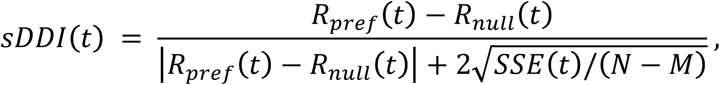

where *t* indicates the time from the stimulus onset. The firing rates were computed over the window ± 5 ms around the time *t*. Note that we defined the preferred and null disparities based on the responses over the entire stimulus duration (as for the standard DDI), because our aim was to quantify the instantaneous selectivity that was congruent to the overall selectivity. If we defined the preferred and null disparities at each time *t*, the sDDI value would always be above zero (due to the circular logic), complicating the interpretation. With our definition, the sDDI could have a negative value when the selectivity at time *t* differed from the selectivity over the entire stimulus duration. This tended to happen more often when *t* was small, and well before the cell’s response latency. During this period, responses are noisy and the sign of the sDDI is stochastic.

#### Cumulative Gaussian function for the sDDI time course

We fitted the cumulative Gaussian function with an amplitude parameter (Figure 8B) to the sDDI time course for MT and V4. During the fitting, we minimized the sum of the squared errors between the fitted function and the mean sDDI (averaged across cells in each area). We used bootstrapping to estimate the 68% confidence intervals of fitted parameter values. When constructing an artificial data set, we resampled the neurons with replacement for each area (N = 83 and 78 for MT and V4, respectively) and computed the new sDDI mean value at each time point. The confidence interval was computed based on 10,000 resampled data sets.

#### Computation of the area ratio without model fitting

We computed the area ratio entirely without model fitting and confirmed the same conclusion. For this model-free analysis, we substituted the amplitude parameter of the fitted Gabor function with the peak-to-trough difference of raw disparity-tuning data. The sign of the amplitude ratio was determined using the sign of the correlation coefficient between the raw tuning data at 100% binocular correlation and those at lower binocular correlations. This was because the tuning-curve inversion should lead to a negative correlation coefficient. We also substituted the quadratic-function fitting to signed amplitude-ratio data with the piecewise linear interpolations between two adjacent raw data points. Then, we calculated the area ratio based on the areas surrounded by these piecewise linear interpolations and the horizontal baseline (zero signed amplitude ratio). The numerator and denominator of the area ratio were the net areas below and above baseline, respectively. For the model-free analysis, we used our entire data set (83 MT and 78 V4 disparity-selective neurons).

## Supplemental Information

**Figure S1. Related to Figure 2C and D.**
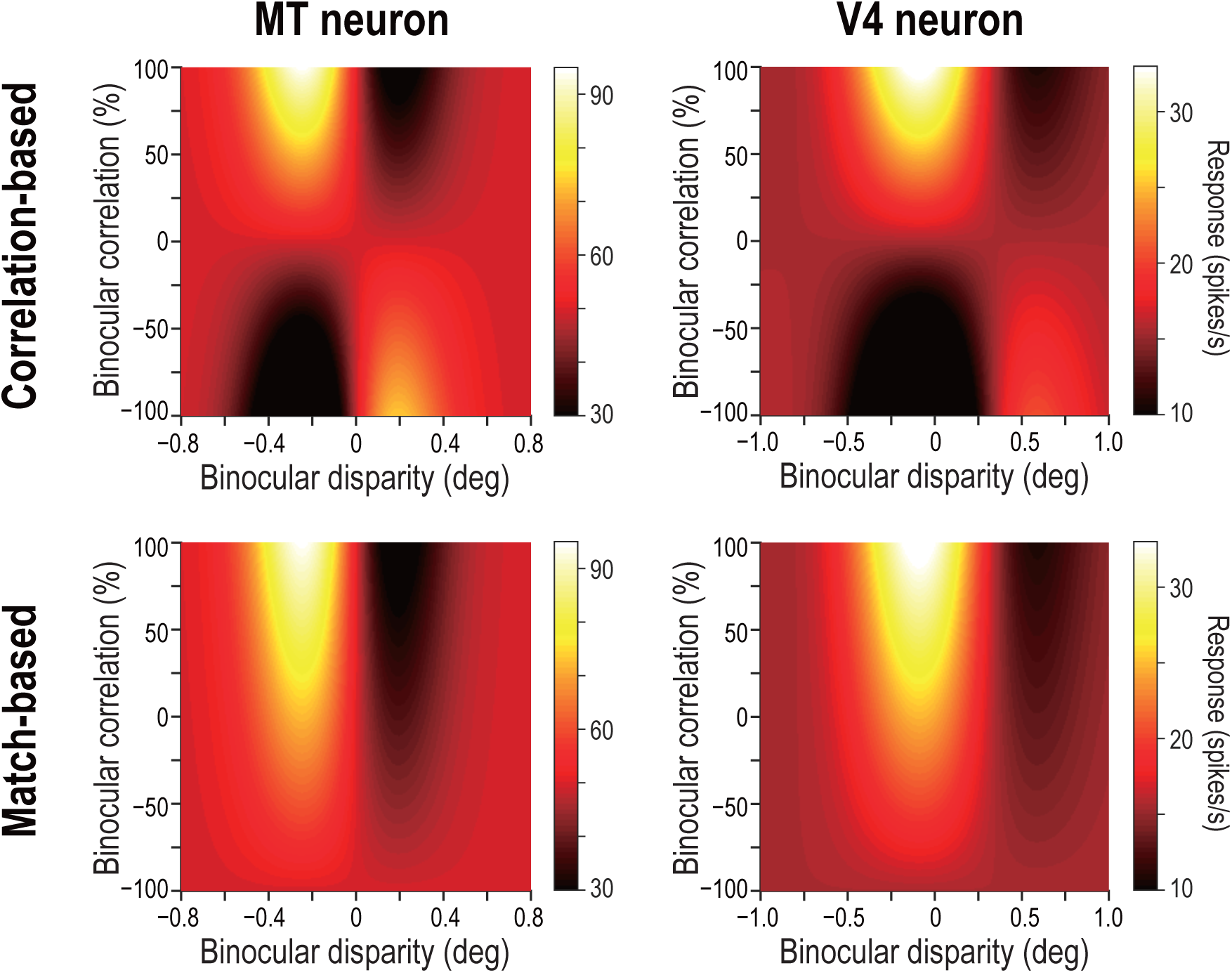
Predicted response maps for example MT and V4 neurons. Compare the observed response maps for the same neurons (shown in Figures. 2C,D) and the predicted maps shown here. The observed map for the example MT neuron is closer to the correlation-based prediction (**top left**), whereas the observed map for the V4 neuron Is closer to the match-based prediction (**bottom right**). We constructed the predictions for each example neuron as follows: we started with the Gabor functions fitted to the disparity-tuning data for correlated RDSs (i.e., at 100% binocular correlation). Next, we computed the predicted responses at lower correlation levels using the linear functions shown In Figure 4A; these functions dictate how the tuning amplitude should change with decreasing binocular correlation for pure correlation-based and match-based models. Specifically, we multiplied the signed amplitude ratio of the correlation-based representation or of the match-based representation by the Gabor function at 100% correlation. The predictions were plotted using the same color ranges as the corresponding observed data.

**Figure S2. Related to Figures 5A-C.**
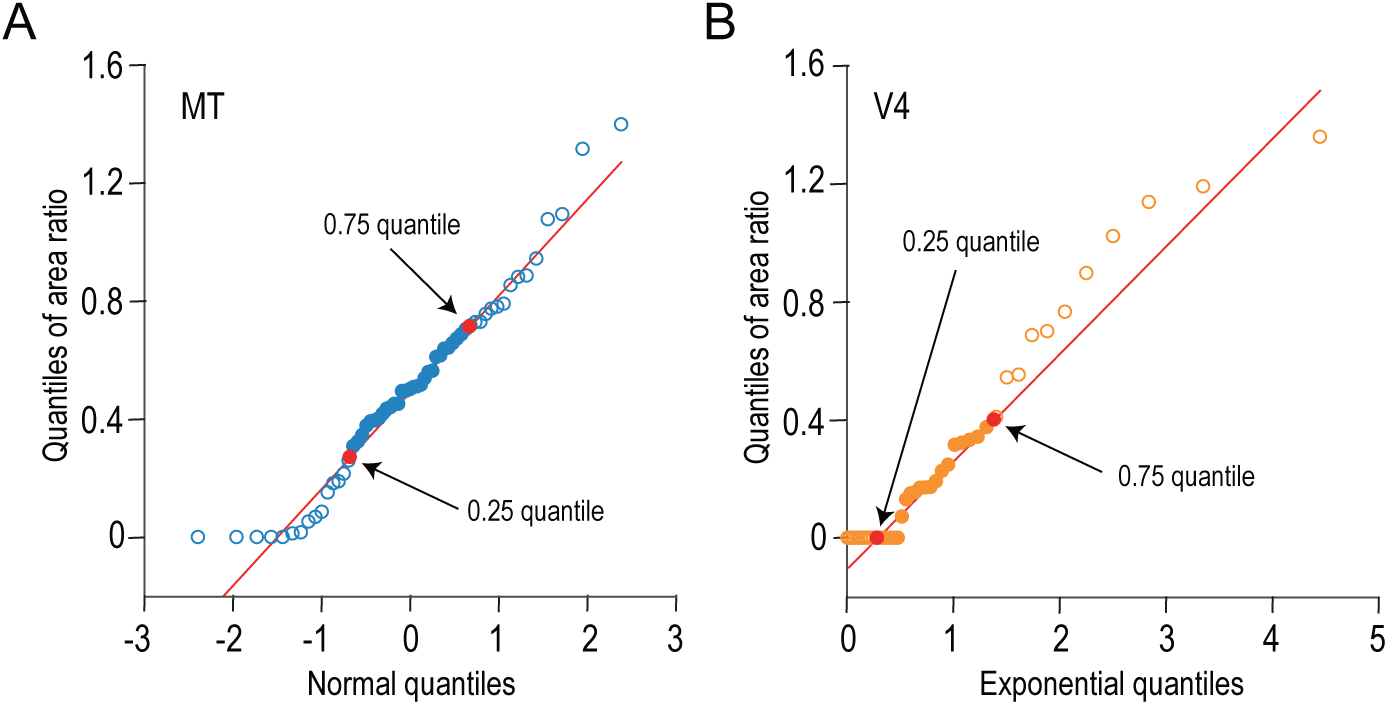
The distributions of the area ratios for MT and V4 are compared against normal and exponential distributions, respectively. We examined the shapes of area ratio distributions using quantile-quantile plots. (**A**) The quantiles of area ratios for MT are plotted against the theoretical quantiles expected from the standard normal distribution. (**B**) Likewise, the quantiles of V4 data are plotted against the theoretical quantiles expected from an exponential distribution (mean = 1). Red lines are drawn through the 0.25 and 0.75 quantiles. The data between these two points are well aligned on the red lines, meaning that the area ratio distributions from MT and V4 follow the normal and exponential distributions, respectively, within the middle 50% of the data range.

**Figure S3. Related to Figure 5C.**
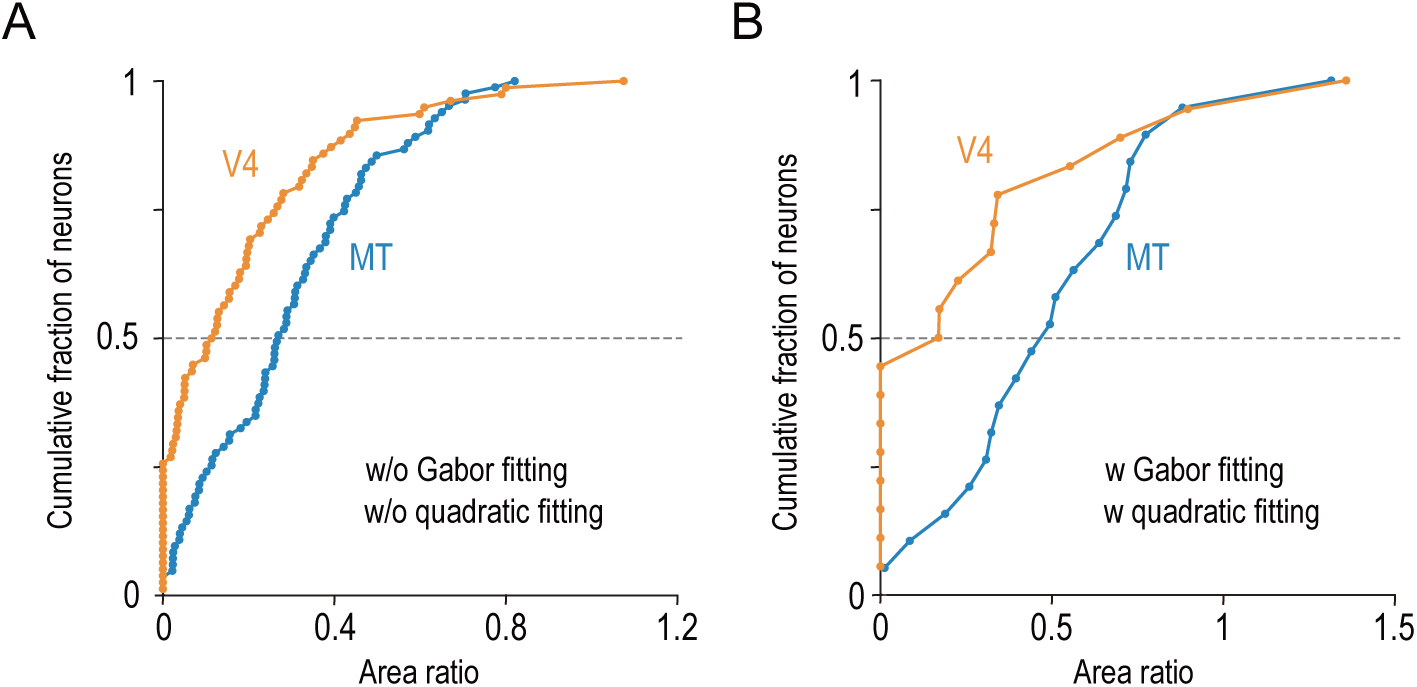
The differential distributions of the area ratios between V4 and MT hold in two control analyses. (**A**) We quantified the area ratio without relying on any model fitting (see Methods). This analysis Included our whole data set (MT, n = 83; V4, n = 78). (**B**), We analyzed only the data obtained from MT and V4 of the same Individual monkey (Monkey 0; MT, n = 28; V4, n = 30).

**Figure S4. Related to Figure 6.**
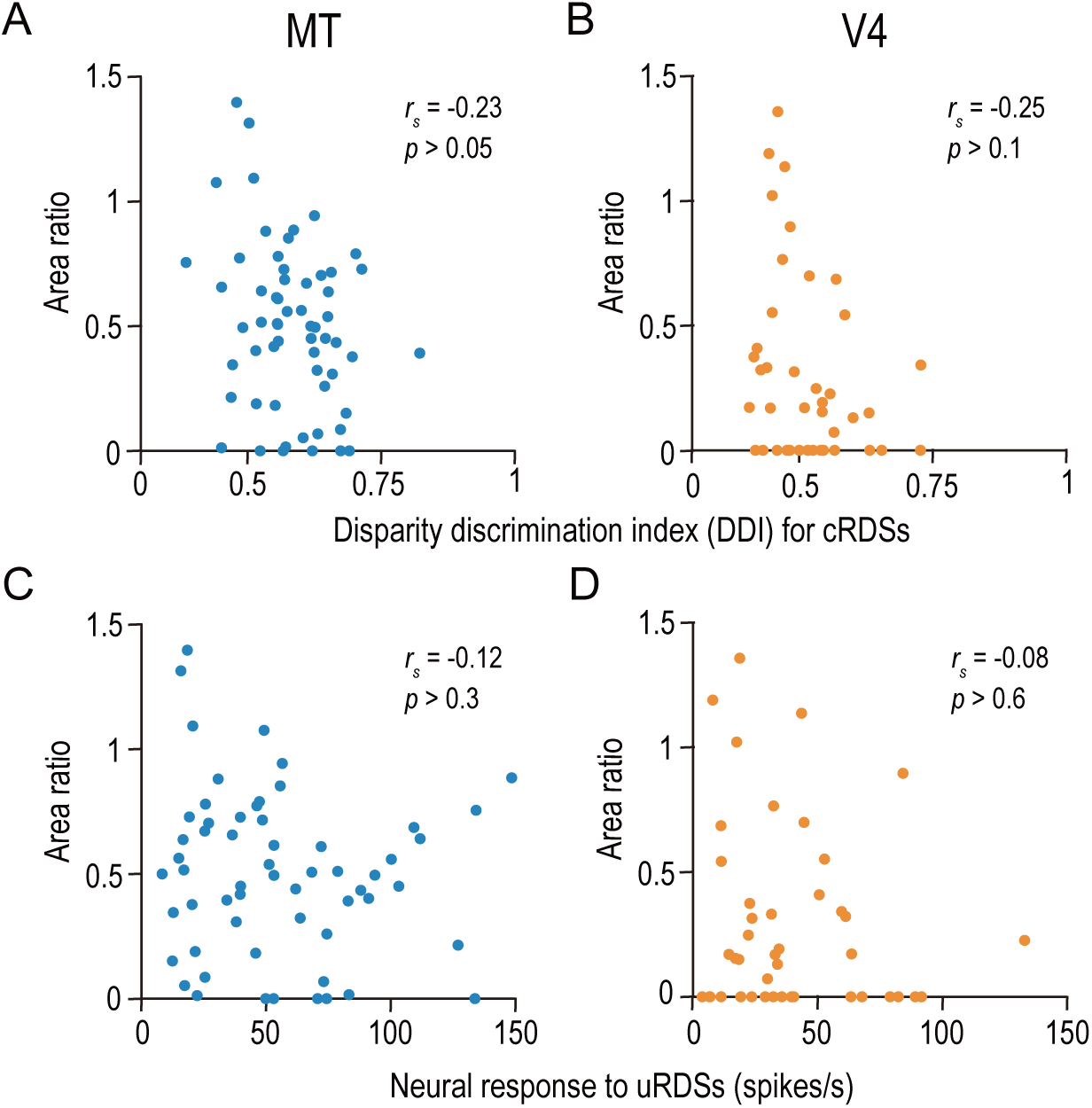
The area ratio is correlated neither with the strength of disparity selectivity nor with the responsiveness In MT and V4. (**A, B**) The area ratio is plotted against the disparity discrimination index (DDI) calculated from the responses to correlated random dot stereograms (cRDSs). The DDI quantifies the strength of disparity selectivity. (**C, D**) The area ratio is plotted against the average firing rate for uncorrelated RDSs, which is often taken as the baseline response level (Uka and DeAngelis, 2003). rs, Spearman’s correlation coefficient. The analyses were performed in the same population of neurons as in Figure 6.

**Figure S5. Related to Figure 6.**
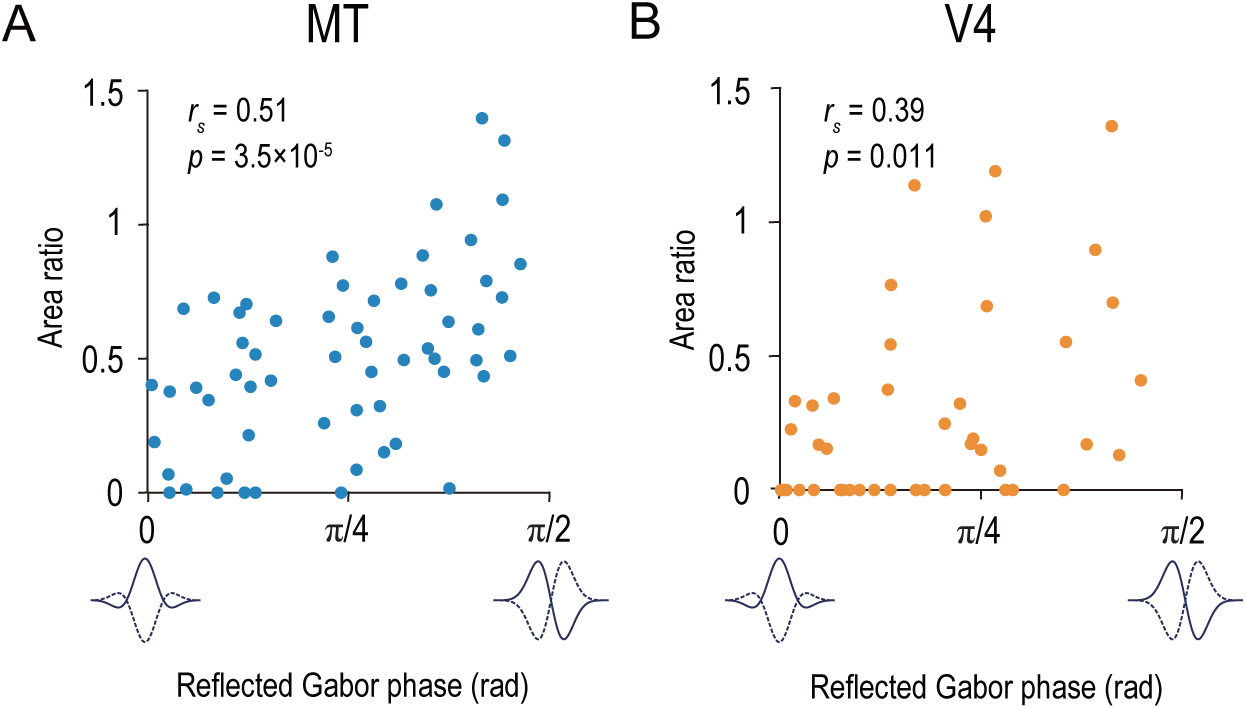
The area ratio is correlated with the symmetry of the disparity-tuning curve as quantified by the Gabor function’s phase parameter in MT and V4. The area ratio is plotted against the best-fitted values of the phase parameter of the Gabor function for 59 MT neurons (**A**) and 43 V4 neurons (**B**). The analyses were performed in the same population of neurons as in Figure 6.

## References

Abdolrahmani, M., Doi, T., Shiozaki, H.M., and Fujita, I. (2016). Pooled, but not single-neuron, responses in macaque V4 represent solution to the stereo correspondence problem. J. Neurophysiol. 115, 1917–1931.

Aoki, S.C., Shiozaki, H.M., and Fujita, I. (2017). A relative frame of reference underlies reversed depth perception in anticorrelated random-dot stereograms. J. Vis. 17, 1–17.

Asher J.M., and Hibbard, P.B. (2018). First- and second-order contributions to depth perception in anti-correlated random dot stereograms. Sci. Rep. 8, 1–19.

Attneave, F. (1954). Some informational aspects of visual perception. Psychol. Rev. 61, 183–193.

Barlow, H.B. (1961). Possible principles underlying the transformations of sensory messages. In Sensory Communication (ed. Rosenblith, W.A.) pp. 217–234. MIT Press, Cambridge, MA.

Borra, E., Belmalih, A., Calzavara, R., Gerbella, M., Murata, A., Rozzi, S., and Luppino, G. (2008). Cortical connections of the macaque anterior intraparietal (AIP) area. Cereb. Cortex. 18, 1094–1111.

Borra, E., Ichinohe, N., Sato, T., Tanifuji, M., and Rockland, K.S. (2010). Cortical connections to area TE in monkey: hybrid modular and distributed organization. Cereb. Cortex 20, 257–270.

Britten, K.H., Shadlen, M.N., Newsome, W.T., and Movshon, J.A. (1993). Response of neurons in macaque MT to stochastic motion signals. Vis. Neuwosci. 10, 1157–1169.

Chen, G., Lu, H.D., Tanigawa, H., and Roe, A.W. (2017). Solving visual correspondence between the two eyes via domain-based population encoding in non-human primates. Proc. Natl. Acad. Sci. USA 114, 13024–13029.

Cheng, K., Hasegawa T., Saleem K.S., and Tanaka, K. (1994). Comparison of neuronal selectivity for stimulus speed, length, and contrast in the prestriate visual cortical areas V4 and MT of the macaque monkey. J. Neurophysiol. 71, 2269–2280.

Chowdhury, S.A., and DeAngelis, G.C. (2008). Fine discrimination training alters the causal contribution of macaque area MT to depth perception. Neuron 60, 367–377.

Cottereau, B.R., Ales, J.M., and Norcia, A.M. (2014). The evolution of a disparity decision in human visual cortex. Neuroimage 92, 193–206.

Cumming, B.G., and DeAngelis, G.C. (2001). The physiology of stereopsis. Annu. Rev. Neurosci. 24, 203–238.

Cumming, B.G., and Parker, A.J. (1997). Responses of primary visual cortical neurons to binocular disparity without depth perception. Nature 389, 280–283.

DeAngelis, G.C., Cumming, B.G., and Newsome, W.T. (1998). Cortical area MT and the perception of stereoscopic depth. Nature 394, 677–680.

DeAngelis, G.C., Ohzawa, I., and Freeman, R.D. (1991). Depth is encoded in the visual cortex by a specialized receptive field structure. Nature 352, 156–159.

DeAngelis, G.C., and Uka, T. (2003). Coding of horizontal disparity and velocity by MT neurons in the alert macaque. J. Neurophysiol. 89, 1094–1111.

Desimone, R., and Schein, S.J. (1987). Visual properties of neurons in area V4 of the macaque: sensitivity to stimulus form. J. Neurophysiol. 57, 835–868.

Dijkerman, H.C., and de Haan E.H.F. (2007). Somatosensory processes subserving perception and action. Behav. Brain Sci. 30, 189–239.

Doi, T., Abdolrahmani, M., and Fujita, I. (2018). Spatial pooling inherent to intrinsic signal optical imaging might cause V2 to resemble a solution to the stereo correspondence problem. Proc. Natl. Acad. Sci. USA 115, 201807687. doi: 10.1073/pnas.1807687115.

Doi, T., and Fujita, I. (2014). Cross-matching: a modified cross-correlation underlying threshold energy model and match-based depth perception. Front. Comput. Neurosci. 8, 1–15.

Doi, T., Takano, M., and Fujita, I. (2013). Temporal channels and disparity computations in stereoscopic depth perception. J. Vis. 13, 1–25.

Doi, T., Tanabe, S., and Fujita, I. (2011). Matching and correlation computations in stereoscopic depth perception. J. Vis. 11, 1–16.

Farivar, R. (2009). Dorsal-ventral integration in object recognition. Brain Res. Rev. 61, 144–153.

Fleet, D.J., Wagner, H., and Heeger, D.J. (1996). Neural encoding of binocular disparity: energy model, position shifts and phase shifts. Vision Res. 36, 1839–1857.

Freud, E., Plaut, D.C., and Behrmann, M. (2016). ‘What’ is happening in the dorsal visual pathway. Trends Cogn. Sci. 20, 773–784.

Fujita, I., and Doi, T. (2016). Weighted parallel contributions of binocular correlation and match signals to conscious perception of depth. Phil. Trans. R. Soc. B 20150257.

Gallyas, F. (1979). Silver staining of myelin by means of physical development. Neurol Res. 1, 203–209.

Gattas, R., and Gross, C.G. (1981). Visual topography of striate projection zone (MT) in posterior superior temporal sulcus of the macaque. J. Neurophysiol. 46, 621–638.

Goncalves, N.R., and Welchman, A.E. (2017). “What not” detectors help the brain see in depth. Curr Biol. 27, 1403–1412.

Goodale, M.A., Milner, A.D. (1992). Separate visual pathways for perception and action. Trends Neurosci. 15, 20–25.

Gu, Y., Watkins, P.V., Angelaki, D.E., and DeAngelis, G.C. (2006). Visual and nonvisual contributions to three-dimensional heading selectivity in the medial superior temporal area. J. Neurosci. 26, 73–85.

Haefner, R.M., and Cumming, B.G. (2008). Adaptation to natural binocular disparities in primate V1 explained by a generalized energy model. Neuron 57, 147–158.

Henriksen, S., Cumming, B.G., and Read, J.C.A. (2016a). A single mechanism can account for human perception of depth in mixed correlation random dot stereograms. PLoS Comput. Biol. 12, e1004906.

Henriksen, S., Read, J.C.A., and Cumming, B.G. (2016b). Neurons in striate cortex signal disparity in half-matched random-dot stereograms. J. Neurosci. 36, 8967–8976.

Hickok, G., and Poeppel, D. (2007). The cortical organization of speech processing. Nat. Rev. Neurosci. 8, 393–402.

Hilgetag, CC. Burns, G.A.P.C., O’Neill, M.A., Scannell, J.W., and Young, M.P. (2000). Anatomical connectivity defines the organization of clusters of cortical areas in the macaque monkey and the cat. Phil. Trans. R. Soc. B 355, 91–110.

Hinkle, D.A., and Connor, C.E. (2002). Three-dimensional orientation tuning in macaque area V4. Nat. Neurosci. 5, 665–670.

Hinkle, D.A., and Connor, C.E. (2005). Quantitative characterization of disparity tuning in ventral pathway area V4. J. Neurophysiol. 94, 2726–2737.

Janssen, P., Verhoef, B.E., and Premereur, E. (2018). Functional interactions between the macaque dorsal and ventral visual pathways during three-dimensional object vision. Cortex 98, 218–227.

Janssen, P., Vogels, R., Liu, Y., and Orban, G.A. (2003). At least at the level of inferior temporal cortex, the stereo correspondence problem is solved. Neuron 37, 693–701.

Julesz, B. (1971). Foundations of Cyclopean Perception (Chicago: Univ Chicago Press).

Krug, K., Cumming, B.G., and Parker, A.J. (2004). Comparing perceptual signals of single V5/MT neurons in two binocular depth tasks. J. Neurophysiol. 92, 1586–1596.

Kumano, H., Tanabe, S., and Fujita, I. (2008). Spatial frequency integration for binocular correspondence in macaque area V4. J. Neurophysiol. 99, 402–408.

Lehky, S.R., and Sereno, A.B. (2007). Comparison of shape encoding in primate dorsal and ventral visual pathways. J. Neurophysiol. 97, 307–319.

Li, P., Zhu, S., Chen, M., Han C., Xu, H., Hu, J., Fang, Y., and Lu, HD. (2013). A motion direction preference map in monkey V4. Neuron 78, 376–388.

Lippert, J., and Wagner, H. (2001). A threshold explains modulation of neural responses to opposite-contrast stereograms. Neuroreport 12, 3205–3208.

Marr, D., and Poggio, T. (1976). Cooperative computation of stereo disparity. Science 194, 283–287.

Marr, D., and Poggio, T. (1979). A computational theory of human stereo vision. Proc. Roy. Soc. Lond. B. Biol. Sci. 204, 301–328.

Masson, G.S., Busettini, C., and Miles, F.A. (1997). Vergence eye movements in response to binocular disparity without depth perception. Nature 389, 283–286.

Mountcastle, V.B., Motter, M.A., Steinmetz, M.A., and Sestokas, A.K. (1987). Common and differential effects of attentive fixation on the excitability of parietal and prestriate (V4) cortical visual neurons in the macaque monkey. J. Neurosci. 7, 2239–2255.

Neri, P. (2005). A stereoscopic look at visual cortex. J. Neurophysiol. 93, 1823–1826.

Nieder, A., and Wagner, H. (2001). Hierarchical processing of horizontal disparity information in the visual forebrain of behaving owls. J. Neurosci. 21, 4514–4522.

Ohzawa, I. (1998). Mechanisms of stereoscopic vision: the disparity energy model. Curr Opin Biolo. 8, 509–515.

Ohzawa, I., DeAngelis, G.C., and Freeman, R.D. (1990). Stereoscopic depth discrimination in the visual cortex: neurons ideally suited as disparity detectors. Science 249, 1037–1041.

Ohzawa, I., DeAngelis, G.C., and Freeman, R.D. (1997). Encoding of binocular disparity by complex cells in the cat’s visual cortex. J. Neurophysiol. 77, 2879–2909.

Orban, G.A., Janssen, P., and Vogels, R. (2006). Extracting 3D structure from disparity. Trends Neurosci. 29, 466–473.

Palanca, B.J.A., and DeAngelis, G.C. (2003). Macaque middle temporal neurons signal depth in the absence of motion. J. Neurosci. 23, 7647–7658.

Parker, A.J. (2007). Binocular depth perception and the cerebral cortex. Nat. Rev. Neurosci. 8, 379–391.

Poggio, G., and Fischer, B. (1977). Binocular interaction and depth sensitivity in striate and prestriate cortex of behaving rhesus monkey. J. Neurophysiol. 40, 1392–1405.

Poggio, G.F., Gonzalez, F., and Krause, F. (1988). Stereoscopic mechanisms in monkey visual cortex: binocular correlation and disparity selectivity. J. Neurosci. 8, 4531–4550.

Prince, S.J.D., Pointon, A.D., Cumming, B.G., and Parker, A.J. (2002). Quantitative analysis of the responses of V1 neurons to horizontal disparity in dynamic random-dot stereograms. J. Neurophysiol. 87, 191–208.

Rauschecker, J.P., and Scott, S.K. (2009). Maps and streams in the auditory cortex: nonhuman primates illuminate human speech processing. Nat. Neurosci. 12, 718–724.

Read, J.C.A., and Cumming, B.G. (2004). Ocular dominance predicts neither strength nor class of disparity selectivity with random-dot stimuli in primate V1. J. Neurophysiol. 91, 1271–1281.

Read, J.C.A., and Cumming, B.G. (2007). Sensors for impossible stimuli may solve the stereo correspondence problem. Nat. Neurosci. 10, 1322–1328.

Roe, A.W., Chelazzi, L., Connor, C.E., Conway, B.R., Fujita, I., Gallant, J.L., Lu, H., and Vanduffel, W. (2012). Toward a unified theory of visual area V4. Neuron 74, 12–29.

Samonds, J.M., Potetz, B.R., Tyler, C.W., and Lee T.S., (2013). Recurrent connectivity can account for the dynamics of disparity processing in V1. J. Neurosci. 33, 2934–2946.

Saur, D., Kreher, B.W., Schnell, S., Kummerer, D., Kellmeyer, P., Vry, MS., Umarova, R., Musso, M., Glauche, V., Abel, S., Huber, W., Rijntjes, M., Hennig, J., and Weiller, C. (2008). Ventral and dorsal pathway for language. Proc. Natl. Acad. Sci. USA 105, 18035–18040.

Seidemann, E., Poirson, A.B., Wandell, B.A., and Newsome, W.T. (1999). Color signals in area MT of the macaque monkey. Neuron 24, 911–917.

Sereno, A.B., and Maunsell, J.H.R. (1998). Shape selectivity in primate lateral intraparietal cortex. Nature 395, 500–503.

Shiozaki, H.M., Tanabe, S., Doi, T., and Fujita, I. (2012). Neural activity in cortical area V4 underlies fine disparity discrimination. J. Neurosci. 32, 3830–3841.

Takemura, A., Inoue, Y., Kawano, K., Qualia, C., and Miles, F.A. (2001). Single-unit activity in cortical area MST associated with disparity-vergence eye movements: evidence for population coding. J. Neurophysiol. 85, 2245–2266.

Tanabe, S., and Cumming, B.G. (2008). Mechanisms underlying the transformation of disparity signals from V1 to V2 in the macaque. J. Neurosci. 28, 11304–11314.

Tanabe, S., Doi, T., Umeda, K., and Fujita, I. (2005). Disparity-tuning characteristics of neuronal responses to dynamic random-dot stereograms in macaque visual area V4. J. Neurophysiol. 94, 2683–2699.

Tanabe, S., Umeda, K., and Fujita, I. (2004). Rejection of false matches for binocular correspondence in macaque visual cortical area V4. J. Neurosci. 24, 8170–8180.

Tanabe, S., Yasuoka, S., and Fujita, I. (2008). Disparity-energy signals in perceived stereoscopic depth. J. Vis. 8, 1–10.

Theys, T., Srivastava, S., van Loon, J., Goffin, J., and Janssen, P. (2012). Selectivity for three-dimensional contours and surfaces in the anterior intraparietal area. J. Neurophysiol. 107, 995–1008.

Uka, T., and DeAngelis, G.C. (2003). Contribution of middle temporal area to coarse depth discrimination: comparison of neuronal and psychophysical sensitivity. J. Neurosci. 23, 3515–3530.

Uka, T., and DeAngelis, G.C. (2004). Contribution of area MT to stereoscopic depth perception: choice-related response modulations reflect task strategy. Neuron 42, 297–310.

Uka, T., and DeAngelis, G.C. (2006). Linking neural representation to function in stereoscopic depth perception: roles of the middle temporal area in coarse versus fine disparity discrimination. J. Neurosci., 25, 6791–6802.

Uka, T., Tanaka H., Kato M., and Fujita I. (1999). Behavioral evidence for visual perception of 3-dimensional surface structures in monkeys. Vision Res., 39, 2399–2410.

Umeda, K., Tanabe, S., and Fujita, I. (2007). Representation of stereoscopic depth based on relative disparity in macaque area V4. J. Neurophysiol. 98, 241–252.

Ungerleider, L.G., and Mishkin, M. (1979). The striate projection zone in the superior temporal sulcus in *Macaca mulatta*: location and topographic organization. J. Comp. Neurol. 188, 347–366.

Ungerleider, L.G., and Mishkin, M. (1982). Two cortical visual systems. In Analysis of Visual Behavior, Ingle, D.J., Goodale M.A., Mansfield, R.J.W. eds. (Cambridge, MA: MIT press), pp. 549–586.

Van Essen, D.C., Maunsell, J.H., and Bixby, J.L. (1981). The middle temporal visual area in the macaque: myeloarchitecture, connections, functional properties, and topographic organization. J. Comp. Neurol. 199, 293–326.

Van Polanen, V., and Davare, M. (2015). Interactions between dorsal and ventral streams for controlling skilled grasp. Neuropsychologia 79, 186–191.

Verhoef, B.E., Vogels, R., and Janssen, P. (2016). Binocular depth processing in the ventral pathway. Phil. Trans. R. Soc. B 20150259.

Wang, Q., Sporns, O., and Burkhalter, A. (2012). Network analysis of corticocortical connections reveals ventral and dorsal processing streams in mouse visual cortex. J. Neurosci. 32, 4386–4399.

Welchman, A.E. (2016). The human brain in depth: how we see in 3D. Annu. Rev. Vis. Sci. 2, 345–376.

Westheimer, G., and Mitchell, A.M. (1956). Eye movement responses to convergence stimuli. A.M.A. Archives Ophthalmology, 55, 848–856.

Yamane, Y., Carlson, E.T., Bowman, J.C., Wang, Z., and Connor, C.E. (2008). A neural code for three-dimensional object shape in macaque inferotemporal cortex. Nature Neurosci. 11, 1352–1360.

